# Behaviorally driven increased glutamatergic activity from lateral entorhinal cortex to hippocampus supports object and spatial memory recall

**DOI:** 10.64898/2026.11.11.765712

**Authors:** M Hernández-Frausto, T Samchok, C D Johnson, O M Bilash, J Tai, J Basu

**Affiliations:** Institute for Translational Neuroscience, New York University Langone Health, New York City, 10016, USA; Department of Neuroscience and Physiology, New York University Grossman School of Medicine, New York City, 10016, USA; Deparment of Psychiatry, New York University Grossman School of Medicine, New York City, 10016, USA; Meinig School of Biomedical Engineering, Cornell University, Ithaca, NY, 14853, USA

## Abstract

Interactions between the entorhinal cortex (EC) and hippocampal area CA1 are vital for episodic memory formation, but their role in recall is less clear. Canonically, spatial processing — the “where” of an experience — is attributed to the medial entorhinal cortex (MEC), whereas the lateral entorhinal cortex (LEC) processes the “what”: non-spatial sensory cues and objects. Though these inputs are anatomically segregated in CA1, multimodal memory processing requires integrating spatial and contextual information, and recent work suggests greater MEC–LEC crosstalk than previously thought. Global LEC silencing via lesions, or combined suppression of glutamatergic and GABAergic LEC neurons, impairs associational goal-oriented learning and multi-object-cued encoding. However, the role of LEC glutamatergic projections specifically to CA1 during recall in freely moving spatial behaviors, and their contribution to synaptic plasticity, remains unclear. We used fiber photometry to record LEC axonal calcium dynamics in CA1 during novel object recognition and Barnes maze tasks, and ex vivo electrophysiology with LEC optogenetic activation. LEC activity increased during recall in both tasks, especially as animals learned spatial cues, peaking at late training stages of the Barnes maze. Local chemogenetic silencing of LEC inputs in CA1 after learning impaired recall. Ex vivo, temporally precise optogenetic activation of LEC input paired with CA3 intrahippocampal inputs was sufficient to induce long-term potentiation of both pathways in CA1 pyramidal neurons. These results suggest that LEC glutamatergic input to CA1 is dynamically modulated by experience to support recall during both object recognition and spatial navigation, with a functional role in synaptic plasticity.

## Introduction

Circuit interactions between the entorhinal cortex and the hippocampus are widely recognized to be crucial for episodic memory^1–3^ Within this framework, the medial entorhinal cortex (MEC) has traditionally been associated with spatial coding through grid, head direction, and border cells^4–8^, whereas the lateral entorhinal cortex (LEC) has typically been described as a “non-spatial” input linked to object- and local cue-related information^9–19^. This has led to the widespread canonical view in which MEC supports spatial mapping by encoding the “where”, while LEC provides mainly content-specific signals about the “what” of the experience^20,21^.

This distinction has become less stark in recent years. Increasing evidence suggests that LEC provides behaviorally relevant variables, including task rules, motivational goals, and salient outcome-predictive cues^19,22–28^. Consistent with this, LEC neurons can track reward- and punishment-related signals, pointing to a role in learning higher-order contextual associations^18,19,21,22,28^ that integrate behaviorally meaningful information. At the same time, lesion and inactivation studies targeting the MEC or LEC have often yielded relatively mild or context-dependent behavioral effects, particularly in standard spatial tasks. This raises an important question: if entorhinal inputs are not strictly required for baseline spatial representations, what is their key contribution to episodic memory? One possibility is that entorhinal inputs, especially from LEC, become more relevant when hippocampal representations need to be associated with context - based rules, stabilized for reliable recall of associations, or linked to behaviorally relevant demands.

Supporting this idea, recent work has shown that LEC inputs can directly influence hippocampal coding dynamics. Disruption of LEC projections to CA3 affects place cell remapping following environmental changes^29^, and more recent work from our group shows that CA3-projecting LEC neurons contribute to goal-oriented learning and the associated stabilization of hippocampal place maps^30^. In addition, recordings of LEC axons within hippocampal circuits indicate that these projections are engaged during goal-directed behavior^19,22^ and may provide information related to object, places and contest needed for episodic memory formation into the hippocampus^14,24^. Together, these findings suggest that LEC inputs do more than convey non-spatial content; they also shape how hippocampal representations relate to behavior. Despite these advances, it remains unclear how LEC input dynamics at their hippocampal targets, specifically in area CA1, are modulated during behavior, and whether these inputs are causally required for memory consolidation, beyond their influence on activity during learning. It remains unknown whether LEC to CA1 projections contribute to the stabilization of contextual and spatial information in the hippocampus over time.

Furthermore, the recent studies suggesting a role for lateral entorhinal cortex in supporting associational learning^19,22,28,30^. However, direct evidence for the role of LEC multimodal inputs in supporting memory recall yet to be explored. We hypothesize that given that LEC helps support stability of spatio-contextual memory representations^30,31^, LEC input to hippocampus is likely important for memory recall beyond its role in learning.

Here, we addressed these questions by combining fiber photometry of LEC axons in dorsal CA1 with pathway- and region-specific chemogenetic silencing in freely behaving mice. We found that LEC to CA1 projections are strongly active during the consolidation phase of novel object recognition, and that their activity increases as animals learn to use global spatial cues in the Barnes maze. Consistent with a causal role, silencing this pathway impaired memory recall in both tasks. Moreover, when using optogenetic activation in ex-vivo slices, we examined ITDP (input timing dependent plasticity) as a striking form of heterosynaptic plasticity. We found that when pairing activity in the CA3 and LEC excitatory axons form a robust information flow through the hippocampal trisynaptic path tuning the efficacy of CA1 pyramidal neurons (PN).

## Results

### Glutamatergic LEC axons in CA1 are more active during the recall phase of the Novel Object Recognition (NOR) task

LEC is involved in processing object-related information, particularly for discriminating between novel and familiar objects^24,32,33^. However, it is unclear exactly when, during behavior, object-related information is active from LEC to the hippocampus and how is modulated during the encoding and recall of object information. To assess the activity dynamics of LEC input to CA1 during the learning and recall phases of object coding behavior, we used fiber photometry-based axonal imaging of LEC glutamatergic projections in CA1 during a freely moving NOR task.

First, we unilaterally expressed the red-shifted genetically encoded Ca^2+^ indicator RgeCO1a-mCherry in LEC glutamatergic neurons (LEC_GLU_) in a Cre-dependent manner using a recombinant adeno-associated virus (rAAV^cre^) in CaMKII-Cre mice. We verified the injection targeting in LEC and projection gradient LEC_GLU_ axons along the proximo-distal axis of dorsal area CA1 (Figure 1a-b). Viral expression was restricted to LEC excitatory neurons (Extended Fig 1A-F) and LEC-CA1 axons exhibited an increasing gradient from the proximal to distal axis in area CA1 (Figure 1C), consistent with the literature. We cross-verified this with independent viral tracing using a CaMKII promoter-driven Chronos-GFP in wild-type animals. Analyzing the density across the area CA1, we observed a lower intensity in the proximal portion of area CA1 compared to distal area CA1 when analyzing the normalized density. With this data, we confirm that LEC_GLU_ projections into HC are differentially segregated across area CA1, a gradient difference between proximo-distal axis.

**Fig. 1.**
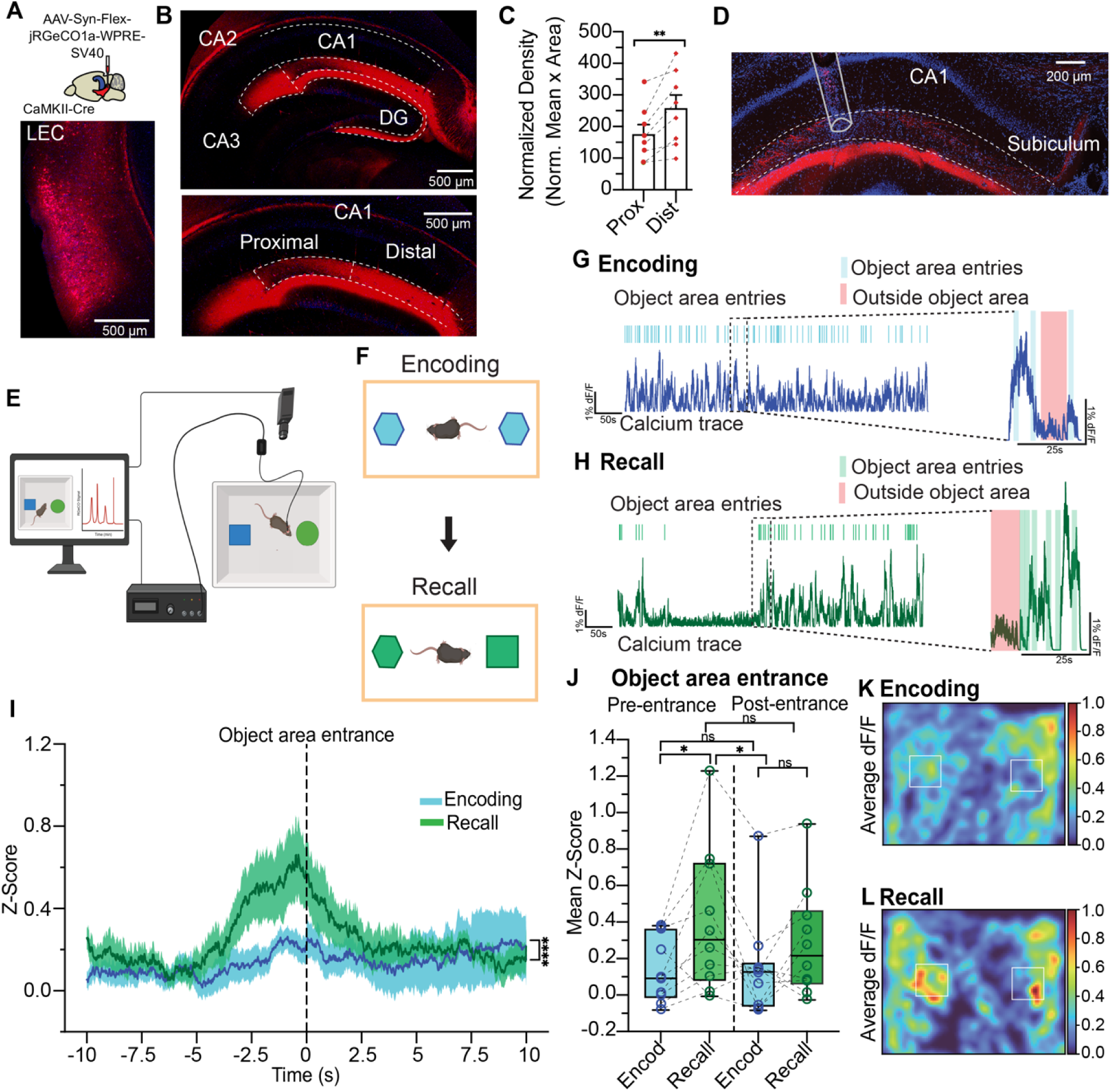
Glutamatergic LEC axons in CA1 are more active during the recall phase of the Novel Object Recognition (NOR) task. **(A)** Schematic representation of the injection strategy for anatomical analysis in LEC (top) and confocal image of the injection site (down). We injected an AAV-SynFLEX-RGECO1a-mCherry in LEC of CAMKII-Cre mice as shown for anatomy analysis. **(B)** Confocal image of coronal brain slice showing the LEC glutamatergic axons in the dorsal hippocampus (red, up) across the different subfields. Confocal image of coronal brain slice showing the proximal and distal proportion of dorsal CA1(red, bottom). **(C)** Quantification of normalized density, analysis is the normalized mean per area, observing a higher density of the glutamatergic projections into the distal portion of area CA1(174 ± 30.40 proximal, 259.5 ± 41.80 distal). Statistical significance was examined with a two-tailed t-test. **p= 0.0018, n= 8. Bar graphs represent columns as the mean with SEM and each dot point represents one slice per animal. **(D)** Confocal image showing glutamatergic axons (in red) from LEC and track of the placement of fiber optic cannulae into distal portion of dorsal hippocampal area CA1, right above SLM layer to record the axonal calcium activity. **(E)** Schematic diagram of the fiber photometry system to record the calcium activity of glutamatergic axons from LEC while mice execute novel object recognition task (NOR). **(F)** Schematic representation of the experimental design of NOR, encoding session is with 2 objects with similar shape, color and size (top) and the recall session with 1 object previously presented and one novel object different in shape and color but similar in size (bottom). **(G)** Representative glutamatergic axonal calcium responses during encoding session (dark blue) highlighting the object area entries (light blue) and outside of object area (light red) signals. **(H)** Representative axonal calcium responses during recall session (dark green) highlighting the object area entries (light green) and outside of the object area (light red) signals. **(I)** Z-score of the average signal during encoding (blue trace) and recall (green trace) sessions of NOR, 10s before the object area entrance (−10s to 0) and 10s after the object area entrance (0 to 10s). Dark lines denote average Z-score, and shaded lines are the mean of all animals, respectively. Statistical significance was evaluated with an ordinary two-way ANOVA. ***p=<0.0001, n=10. **(J)** Mean Z-score of 5s before object area entrance and 5s after object area entrance in encoding (blue) and recall (green) presented as box-and whisker plots (median, lower/upper quartiles and 10-90th percentiles) 5s previous object entrance: mean Z score ± SEM: 0.1411 ± 0.003 encoding, 0.4024 ± 0.009 recall; 5s after object entrance: mean Z score ± SEM: 0.1518 ± 0.002, encoding, 0.2858 ± 0.005 recall. Each data point represents one animal. Statistical significance is an ordinary two-way ANOVA with a significance effect between encoding and recall 5s before object entrance (*p=0.0192) and between recall 5s before entrance and 5s after in encoding (*p=0.0260) sessions. **(K)** Representative heatmap of one mouse showing the averaged and normalized dF/F relative to the objects area zone during the encoding session. **(L)** Representative heatmap of one mouse showing the averaged and normalized dF/F relative to the objects area zone during the recall session. Data are present as mean ± SEM.

Next, we performed *in vivo* fiber photometry recordings during freely moving behavior. We implanted a fiber-optic cannula unilaterally over the distal portion of dorsal *stratum lacunosom moleculare* (SLM) to track behaviorally driven population dynamics of LEC axons in area CA1 (Figure 1D). We imaged axonal calcium dynamics during the encoding and recall phases of the novel object recognition task (NOR, Figure 1E-F). We observed greater axon activity in the object entry zone during recall compared to encoding (Figure 1i-j). This was aligned with a higher amplitude in the pre-entrance of the object zone but not for the AUC (Extended Figure 2A-C). Lastly, we observed a higher correlation activity during recall in the object area zone compared to encoding (Figure 1K-L). These findings suggest that LEC glutamatergic axons in CA1 play an active role specifically during recall in the novel object recognition task.

### Silencing of glutamatergic LEC to CA1 projections during recall impairs novelty recognition in the novel object recognition test

Next, we examined whether the LEC glutamatergic (LEC_GLU_) input to CA1 is necessary for recall of NOR. To test this, we employed a loss-of-function approach of selective chemogenetic silencing of LEC_GLU_ axon terminals in CA1 during behavior. Here, we bilaterally expressed the pharmacogenetically specific actuator module (PSAM4-GlyR)^34^ in LEC glutamatergic neurons using a CaMKII promoter-driven rAAV (Figure 2A). To selectively silence LEC_GLU_ locally in CA1, we implanted a custom-designed bilateral cannula guide over dorsal CA1 to infuse the Pharmacologically Selective Effector Molecule (PSEM817) ligand into the SLM layer. This allowed silencing of LEC_GLU_ axons locally with spatial precision into dorsal CA1, 5 minutes before the recall session of NOR (Figure 2B-D). Our control group consisted in the same PSAM4-GlyR mice but were infused with saline only, and a subset of control mice consisted of a GFP expressing neuron that received PSEM infusion.

**Fig 2.**
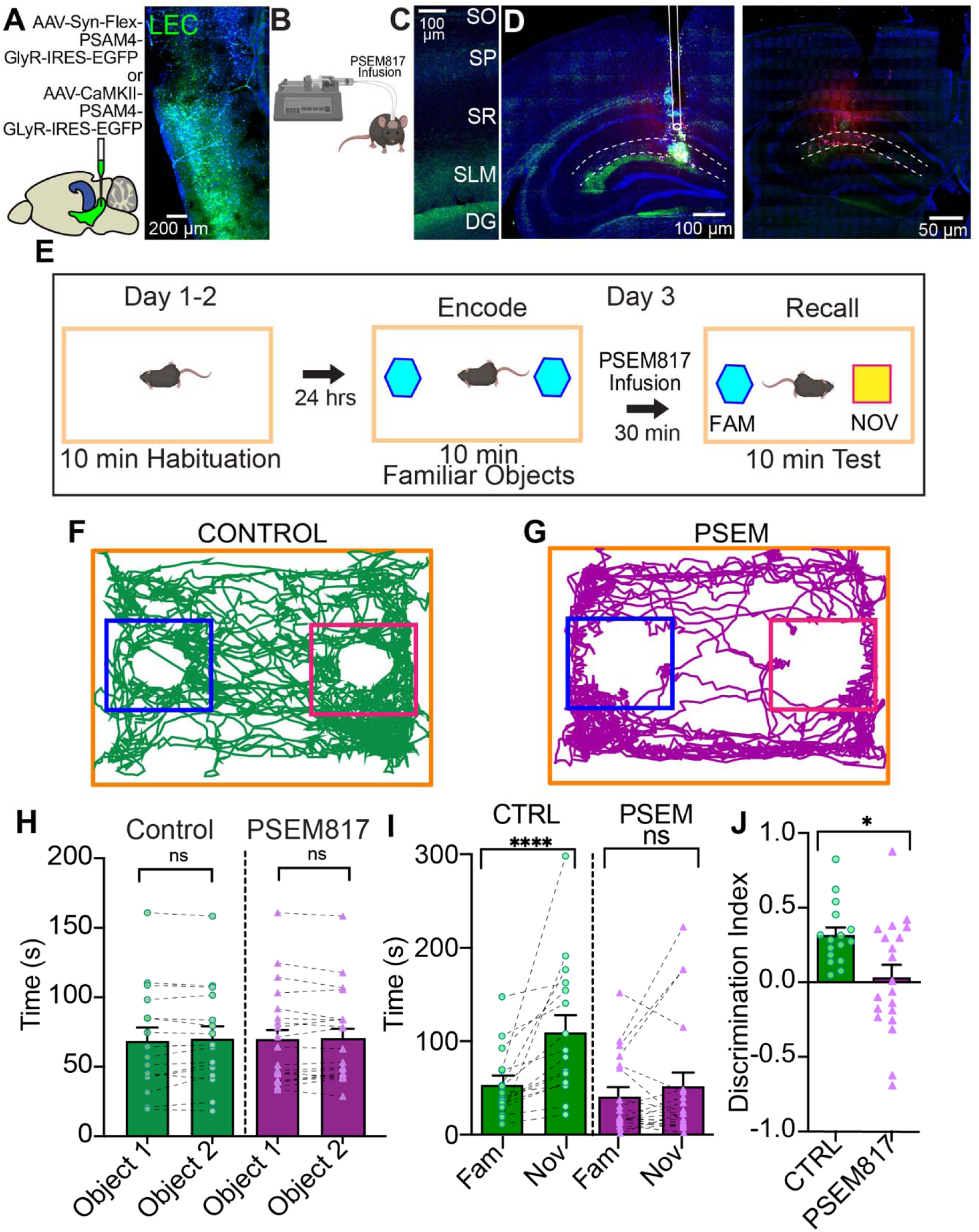
Silencing of glutamatergic LEC to CA1 projections during recall impairs novelty recognition in the novel object recognition test. **(A)** Schematic representation of the injection site for the chemogenetic strategy in LEC (left) and confocal image of the injection site (right). We injected AAV-SynFlex-PSAM4-GlyR-IRES-EGFP or AAV-CaMKII-PSAM4-Gly-IRES-EGFP in CaMKII-cre mice in LEC. **(B)** Schematic representation of chemogenetic silencing strategy of local infusion of PSEM817 ligand to locally silence LEC glutamatergic projections into dorsal area CA1 before test/recall session. **(C)** Confocal image of area CA1 with the glutamatergic axons in SLM (green). **(D)** Confocal image of coronal brain slice showing the hippocampal region with the glutamatergic axons from LEC (green) and the track of the cannula guide for local infusion. Confocal image of coronal brain slice with the hippocampal region showing the restriction of the local infusion of PSEM817 when coupled to a miniRuby (red) fluorophore in hippocampal area CA1 only. **(E)** Experimental design of NOR task. Habituation (day 1-2), mice were placed in an open field arena without any object and allowed to explore for 10 min each day. On day 3, encoding and recall of NOR was evaluated. Here, mice were placed in the arena with two objects similar in shape, size and color and were allowed to explore and familiarize with the objects for 10min. Afterwards, mice were connected to a infusion pump to locally infuse bilaterally the ligand PSEM817 in a 0.5 µl volume (15 µM) or 0.9% saline only into dorsal CA1. After 30 min of the encoding session, the mice were located into the arena but one of the objects was replaced with a novel and different object. Animals were allowed to freely explore the objects for 10 min. **(F)** Representative map of the head trajectory from one mouse in control conditions (locally infused bilaterally of saline into dorsal area CA1),, showing the head distance trajectory and the occupancy (green) into the object area zone from familiar object (blue square) and novel object (red square). **(G)** Representative map of the head trajectory from one mouse in PSEM817 conditions (locally infused bilaterally of PSEM817 into dorsal area CA1), showing the head distance trajectory and the occupancy (dark purple) into the object area zone from familiar object (blue square) and novel object (red square). **(H)** Analysis of the time spent exploring object 1 and object 2 during encoding trial of NOR in control and PSEM817 with no infusion. For control: 53.27 ± 9.061 familiar, 109.4 ± 18.33 novel; for PSEM817: 40.63 ± 9.159 familiar, 51.75 ± 14.68 novel. Statistical significance tested with a two-wailed unpaired U Mann-Whitney test for both cohorts (control p=0.9556; PSEM817 p=0.8464). **(I)** Analysis of the comparison of time that mice spent exploring the novel and the familiar objects in the test trial of NOR task in control and PSEM817 conditions that were locally infused into hippocampal area CA1. Statistical significance was tested with a two-tailed Wilcoxon pairs signed-rank test for both cohorts (***p= < 0.0001 for control, p= 0.6215 for PSEM817). **(J)** Comparison of the novel object discrimination index of mice during NOR test trial. Statistical significance tested with a two-tailed unpaired t-test confirming a significant effect of local infusion silencing with PSEM817 (*p=0.0115). Data are present as mean ± SEM.

Mice were trained in the standard NOR as previously described, and we locally infused the PSEM817 ligand or saline before the recall phase (Figure 2E). Representative locomotor trajectories during recall session showed exploration patterns with occupancy of the familiar and novel object area showing a higher occupancy in novel object area in control rather than PSEM817-infusion animals (Figure 2F-G). When analyzing the amount of time exploring each object during the encoding phase, both groups spent similar amounts of time exploring each object (Figure 2H). During the recall session, control animals displayed a robust preference for the novel object area, whereas PSEM817-treated animals failed to discriminate between familiar and novel objects (Figure 2I), without differences in the distance travelled across the session (Extended Figure 3A-B). Accordingly, the discrimination index was significantly reduced following chemogenetic silencing of LEC_GLU_ terminals in CA1 (Figure 2J), consistent with a decreased NOR index (extended Figure 3C). Interestingly, this effect was not observed when the silencing of LEC_GLU_ axon terminals in CA1 were done before encoding phase of NOR (extended Figure 4A-H). Together, these results indicate that LEC_GLU_ inputs to CA1 are contribute for memory retrieval during NOR.

### Glutamatergic LEC-CA1 inputs show increased activity as mice learn the escape hole location in the Barnes maze

Recent studies have questioned a strict functional segregation of the LEC as exclusively involved in contextual processing, instead highlighting substantial crosstalk in spatial coding with MEC and suggesting a broader contribution of LEC to spatially guided behavior ^35–37^. Thus, we sought to dissect the functional role of LEC glutamatergic inputs (LEC_GLU_) to dorsal hippocampal area CA1 during learning and recall in a spatial navigation task using the Barnes maze, by evaluating the population activity of glutamatergic axons through fiber photometry (Figure 3A). First, mice were trained for 8 days to locate the escape hole using global distal cues, with 4 trials per day on days 1 to 7 and a single recall trial on day 8 (Figure 3B). Photometry recordings were grouped into 4 performance stages covering early learning to recall. To enable direct comparisons across stages, we analyzed two trials per day for each stage, For the first 3 stages, these corresponded to the first two trials, whereas for stage 4, the analyzed trials were the last from day 7 and the only trial recorded on day 8. (Figure 3I). Axonal calcium activity was aligned to escape hole approaches (Figure 3C-F). Interestingly, we observed an increased Z-score dF/F calcium axonal activity specifically for the escape hole as mice progress through the 4 stages, culminating with a calcium signal highest in stage 4 (Figure 3G). When evaluating the dF/F across the 4 stages and observed a higher average activity in the zone surrounding the escape hole zone once the mice had learned its location (Figure 3H). Finally, while analyzing the specificity of the effect, we observed that mean Z-score was higher in the stage 4 compared to the first 2 stages and once the mice learned and consolidate the location of the escape hole (Figure 3I). Furthermore, other metrics of activity including, amplitude and AUC increased across stages specifically, for the escape hole whereas non escape hole activity remained constant irrespective of the training stage. Notably across animals, events were significantly larger during their interactions with the escape hole as compared to non-escape holes at later stages of learning and recall (stages 3 and 4, extended Figure 5A-C). Together, these results suggest that LEC_GLU_ to dorsal CA1 become highly active as the hippocampus consolidates the location of the escape hole during the Barnes maze task.

**Fig 3.**
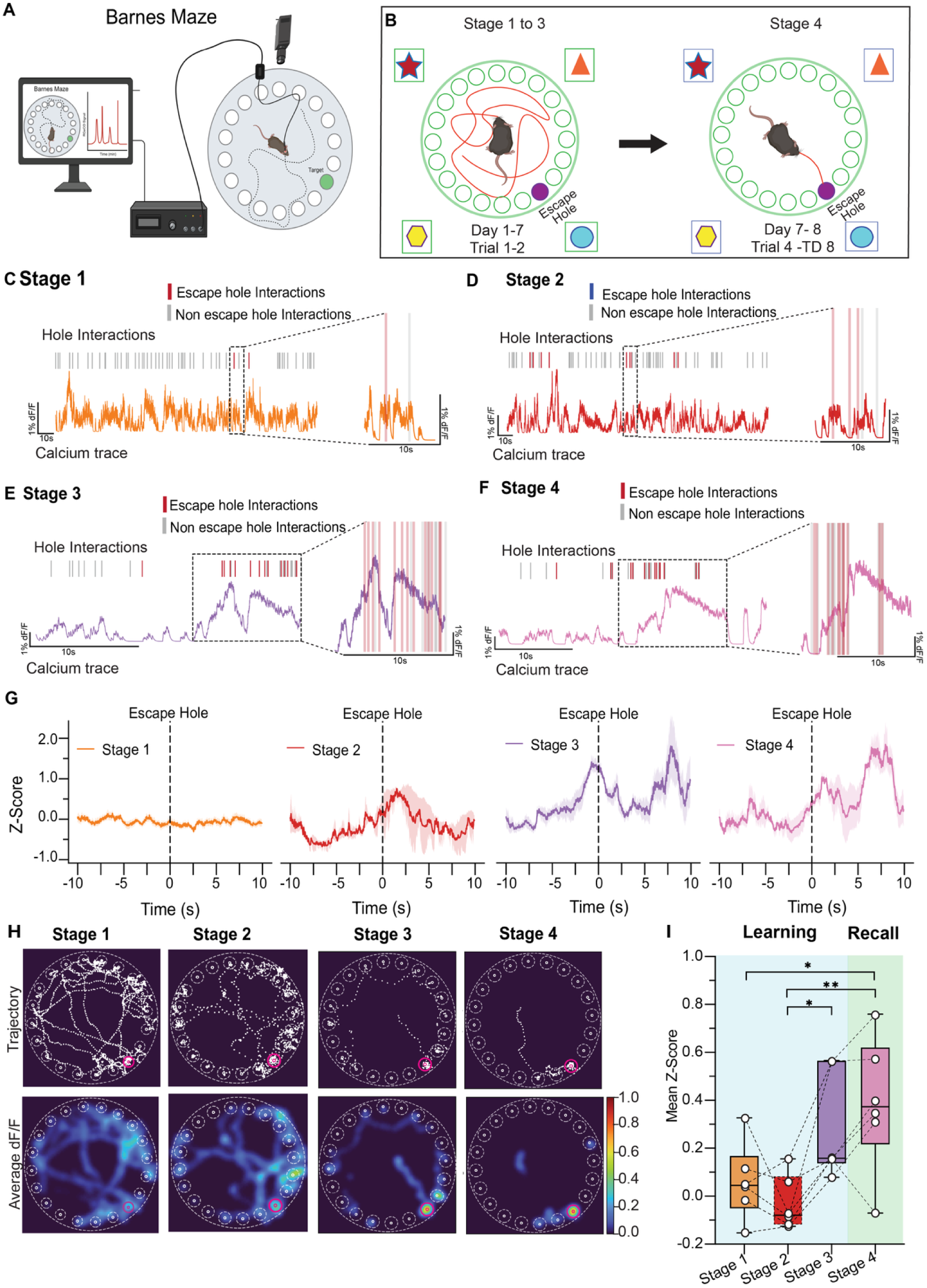
Increased activity of LEC glutamatergic inputs into area CA1 while mice consolidate the location of escape hole during Barnes maze task. **(A)** Schematic diagram of the fiber photometry system to record the calcium activity of glutamatergic axons from LEC into hippocampal dorsal CA1 while mice execute Barnes Maze task. **(B)** Experimental design of Barnes maze task. In the stages 1 to 3 (day 1-7, 2 trials) mice are allowed to freely explore the maze and its holes for 3 minutes or until it finds the escape hole, recordings of the LEC glutamatergic axons are assessed. In the stage 4, mice are allowed to explore the maze until it finds the escape hole. This is the last trial of day 7 and recall test phase (day 8). **(C)** Representative axonal calcium responses during stage 1 (orange) highlighting the escape hole interactions (blue) and non-escape hole interactions (gray). **(D)** Representative axonal calcium responses during stage 2 (red) highlighting the escape hole interactions (blue) and non-escape hole interactions (gray). **(E)** Representative axonal calcium responses during stage 3 (purple) highlighting the escape hole interactions (blue) and non-escape hole interactions (gray). **(F)** Representative axonal calcium responses during stage 4 (pink) highlighting the escape hole interactions (blue) and non-escape hole interactions (gray). **(G)** Z-score of the average signal during the 4 stages of Barnes maze task, for stage 1 (orange trace), for stage 2 (red trace), for stage 3 (purple trace), and for stage 4 (pink trace). Z-score is from 10s before the escape hole interaction (−10 to 0s) and 10s after the escape hole interaction (0 to 10s). Dark lines denote average Z-score, and shaded lines are the mean of all animals, respectively. **(H)** Representative heatmap of one mouse showing the trajectory (top graph) and averaged normalized dF/F relative to escape hole interactions during stages (bottom graph) from stage 1 to 4. **(I)** Mean Z-score of 10s after escape hole interactions during the learning (light blue box) for stage 1 (orange), stage 2 (red) and stage 3 (purple) and in recall for stage 4 (pink) presented as a box-and whisker plots (median, lower/upper quartiles and 10-90th percentiles) (0.006 ± 0.045 stage 1, 0.060 ± 0.1324 stage 2, 0.305 ± 0.107 stage 3, 0.4781 ± 0.083 stage 4). Each data point represents one animal. Statistical significance is an ordinary two-way ANOVA with a significance between stage 1 and 4 (*p= 0.0152) and between stage 2 and 4 (*p= 0.0315). Data are present as mean ± SEM.

### Glutamatergic LEC-CA1 inputs are required for spatial memory retrieval in the Barnes maze task

Because we observed that LEC glutamatergic inputs (LEC_GLU_) to CA1 are highly active while mice consolidate and recall the location of the escape hole in the Barnes maze task, we next asked whether these projections contributed to spatial memory recall in this task. To address this question, we used the same chemogenetic strategy of PSAM4-GlyR in CAMKII promoter as in our previous experiments (Figure 2A-D), with bilateral cannulae above dorsal CA1 to locally infuse PSEM817 or saline 5 minutes before recall session of the Barnes maze paradigm (Figure 4A-B). All mice were first trained to locate the escape hole using distal spatial cues across seven days of learning sessions consisting of four trials per day, followed by a single recall trial (Figure 4B).

**Fig 4.**
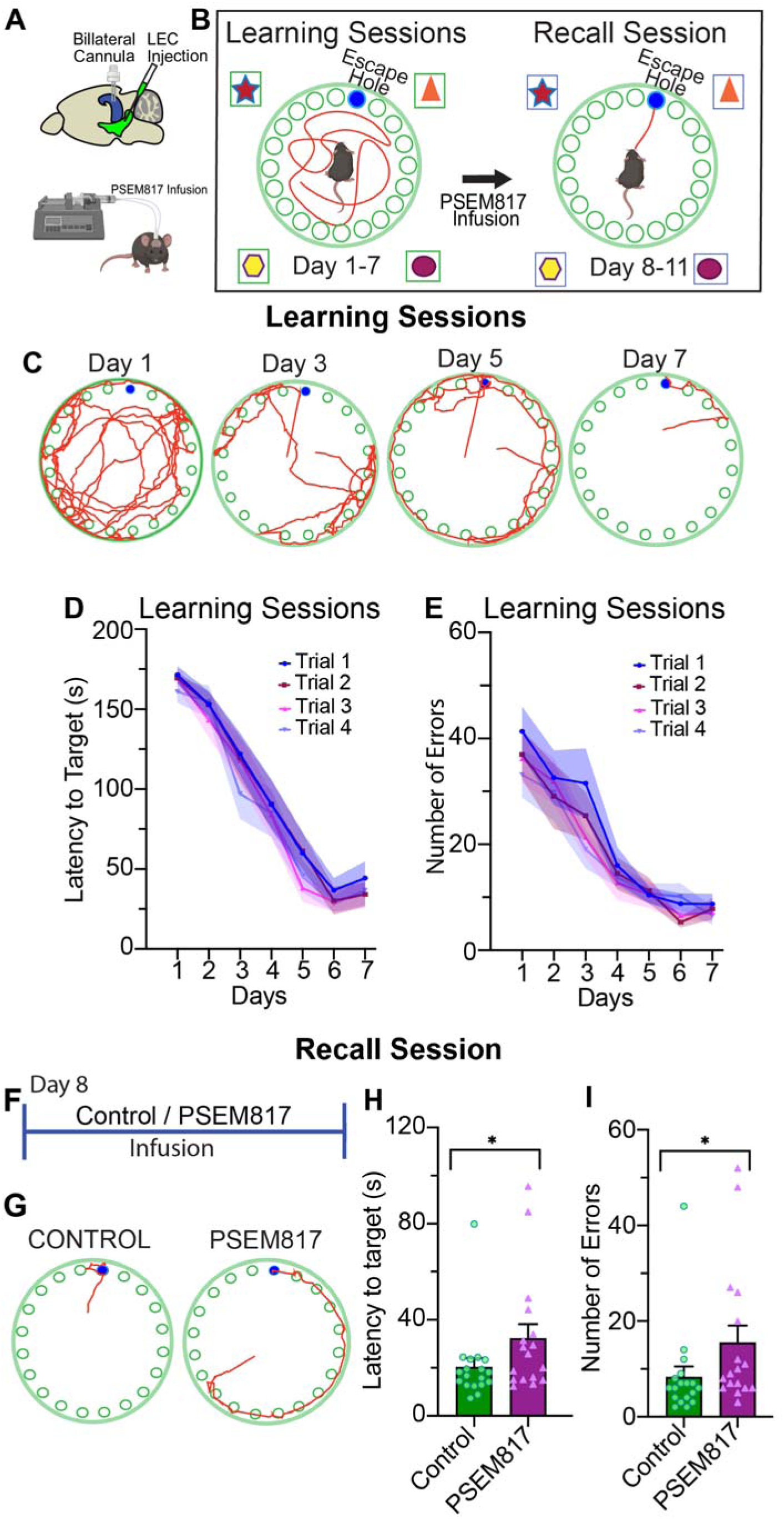
LEC Glutamatergic inputs into dorsal CA1 are necessary for spatial consolidation in the Barnes maze task. **(A)** Schematic representation of chemogenetic silencing strategy with injection in LEC and cannulae place into dorsal area CA1 for local infusion of PSEM817 ligand to locally silence LEC glutamatergic projections into dorsal area CA1. **(B)** Schematic representation of experimental design for Barnes maze task. In learning sessions (day 1-7) mice were placed in the escape box for 1 min. Then, mice were placed at the center of the maze and allowed to freely explore the maze and its holes for 180s (3 min) or until it finds the escape hole, 4 trials per day were assessed. In the recall session, mice were allowed to explore the maze until it finds the escape hole. Before the recall session, infusion into dorsal CA1 of saline or PSEM817 only were delivered for control and PSEM conditions. **(C)** Representative map of the head trajectory from one mouse during learning sessions on day 1, day 3, day 5 and day 7. **(D)** Latency to target during learning sessions for 7 days in trial 1 (blue line), trial 2 (dark red), trial 3 (pink), trial 4 (purple). For analyzing the differences between trials, statistical significance was obtained by fitting a full-model two-way ANOVA, and no differences were observed among trials in each day (167.7 ± 2.307 day 1, 150.9 ± 2.625 day 2, 114.1 ± 5.737 day 3, 87.39 ± 1.792 day 4, 51.11 ± 5.667 day 5, 31.11 ± 1.921 day 6, 37.32 ± 2.403 day 7). For analysis of differences across learning session days, statistical significance was obtained by an ordinary two-way ANOVA, confirming a significant effect between day 1 and day 3,4,5,6,7 (***p=<0.0001) but not day 2, day 2 and day 3,4,5,6,7 (***p=<0.0001) but not day 1, day 3 and day 1,2,4,5,6,7 (***p= 0.0014 day 3 versus day 4, ***p=<<0.0001 day 5,6,7), day 4 and day 1,2,3,5,6,7 (***p=<<0.0001), day 5 and day 1,2,3,4,6 but not 7 (***p=<<0.0001 day 1,2,3,4, *p= 0.0452 day 6), day 6 and day 1, 2, 3, 4, 5 but not 7 (***p=<0.0001 day 1,2,3,4, *p= 0.0452 day 5), day 7 and day 1,2,3,4,5 but not 6 (***p=<0.0001). **(E)** Number of errors to enter escape hole during learning sessions for 7 days in trial 1 (blue line), trial 2 (dark red), trial 3 (pink), trial 4 (purple). For analyzing the differences between trials, statistical significance was obtained by fitting a full-model two-way ANOVA, and no differences were observed among trials in each day (36.89 ± 1.692 day 1, 30.82 ± 0.831 day 2, 24.31 ± 2.739 day 3, 14.24 ± 0.689 day 4, 10.71 ± 0.187 day 5, 7.679 ± 1.079 day 6, 7.536 ± 0.472 day 7). For analysis of differences across learning session days, statistical significance was obtained by an ordinary two-way ANOVA, confirming a significant effect between day 1 and day 3,4,5,6,7 (***p=<0.0001) but not day 2, day 2 and day 3,4,5,6,7 (*p= 0.0476 day 2 versus day 3, ***p=<0.0001 day 4,5,6,7), day 3 and day 1,2,4,5,6,7 (*p=0.0476 day 3 versus day 2, ***p=<0.0001 day 4,5,6,7), day 4 and day 1,2,3,5,6,7 (P<0.0001 day 4 versus day 1,2,3, *p=0.0421 day 4 versus day 6, *p=0.0347 day 4 versus day 7) but not day 7, day 5 and day 1,2,3,4 (***p=<0.0001) but not day 6 and day 7, day 6 and day 1,2,3,4 (***p=<0.0001) but not day 5 and day 7, day 7 and day 1,2,3,4 (***p=<0.0001) but not day 7. **(F)** Timeline of the experimental design of recall session. For local silence in control and PSEM817. **(G)** Representative map of the head trajectory of one mouse in control conditions and with PSEM817 locally infused of saline or PSEM817 into dorsal area CA1, showing the head distance trajectory (red) and the location of the escape hole (blue circle). **(H)** Analysis of the latency to target by mice counted as the total period a mouse explores the arena until entering the escape box during recall sessions (20.31 ± 3.699 control, 32.24 ± 5.913 PSEM817). Statistical significance tested with a two-wailed unpaired U Mann-Whitney test (*p= 0.0413). **(I)** Analysis of the total number of errors made by mice counted as the number of times it explores different holes including the escape hole without entering the escape box during recall sessions (8.278 ± 2.233 control, 15.47 ± 3.584 PSEM817). Statistical significance tested with a two-wailed unpaired U Mann-Whitney test (*p= 0.0292). Data are present as mean ± SEM.

During training, animals progressively improved their performance, as reflected by more direct trajectories toward the escape hole across learning days (Figure 4C). Consistent with this improvement, both latencies to reach the escape hole and the number of errors decreased across learning sessions and trials, indicating successful acquisition of the task (Figure 4D-E). To determine the role of LEC_GLU_ inputs during memory recall, PSEM817 or saline only were infused prior the recall session (Figure 4F). No differences were observed in the distance to target and the mean speed during learning days (extended figure 6A-C). Interestingly, representative trajectories revealed that navigation toward the escape hole, from mice when receiving the infusion of PSEM817, displayed a less direct research strategy (Figure 4G). Consistent with these observations, chemogenetic silencing of the inputs significantly increased the latency required to locate the escape hole during the recall session (Figure 4H). A similar effect was observed when analyzing the number of errors, which was also increased following the silencing of LEC_GLU_ inputs to CA1 (Figure 4I). Together, these findings indicate that LEC_GLU_ inputs to dorsal CA1 contribute to spatial memory recall in the Barnes maze task.

### Glutamatergic LEC-CA1 inputs coupled with CA3 are necessary for ITDP expression

Given our findings that glutamatergic LEC (LEC_GLU_) inputs to CA1 support behavioral recall, we explored the cellular mechanisms of this circuit in memory storage using *ex vivo* patch-clamp electrophysiology and optogenetic activation of LEC_GLU_. Previous studies have highlighted a physiologically relevant non-Hebbian plasticity rule input timing dependent plasticity (ITDP)^25,38–40^, requiring instructive signals from the entorhinal cortex to induce long-term heterosynaptic plasticity of CA3-CA1 Schaffer collateral evoked output in CA1 pyramidal neurons. Typically, ITDP is induced with inhibition intact, using coordinated activity matched in timing to information flow in the cortico-hippocampal delay line circuit (20 ms), and at low frequencies mimicking the activity CA1 neurons during free exploration in rodents (1 Hz). We asked whether LEC_GLU_ inputs may also contribute to the induction of robust, longer-lasting forms of synaptic plasticity (such as ITDP), that may contribute to the increased LEC_GLU_ activity we observed in the later phases of Barnes maze and persistent through memory recall. To test this, we expressed the blue light gated opsin ChR2 in LEC_GLU_ neurons driven by the CaMKII promoter^41^ and activated the LEC_GLU_ inputs with photostimulation in CA1 SLM (Figure 5A-B). Pairing the LEC_GLU_ with electrical stimulation of CA3 inputs was sufficient to induce ITDP in CA1 PNs (Figure 5C). Moreover, we also observed a homosynaptic long-term potentiation (LTP) of the LEC_GLU_ inputs themselves (Figure 5D). Consistent with these findings, we observed that peak PSP amplitudes significantly increased in both CA3 and LEC_GLU_ inputs following ITDP induction and compared with the baseline (Figure 5E). However, when analyzing the PSP amplitudes relative to a theoretical mean of 1, we observed significant potentiation only on CA3-CA1 inputs (Figure 5F). Together, these findings suggest that appropriately timed activation of LEC_GLU_ and CA3 inputs induces a robust and persistent potentiation of both pathways in CA1, with a higher enhancement of CA3-CA1 synaptic transmission. This suggests that LEC_GLU_ can facilitate the strengthening of convergent CA3 inputs through ITDP, providing a potential cellular mechanism by which cortical information modulates hippocampal circuitry activation during memory consolidation and recall.

**Fig 5.**
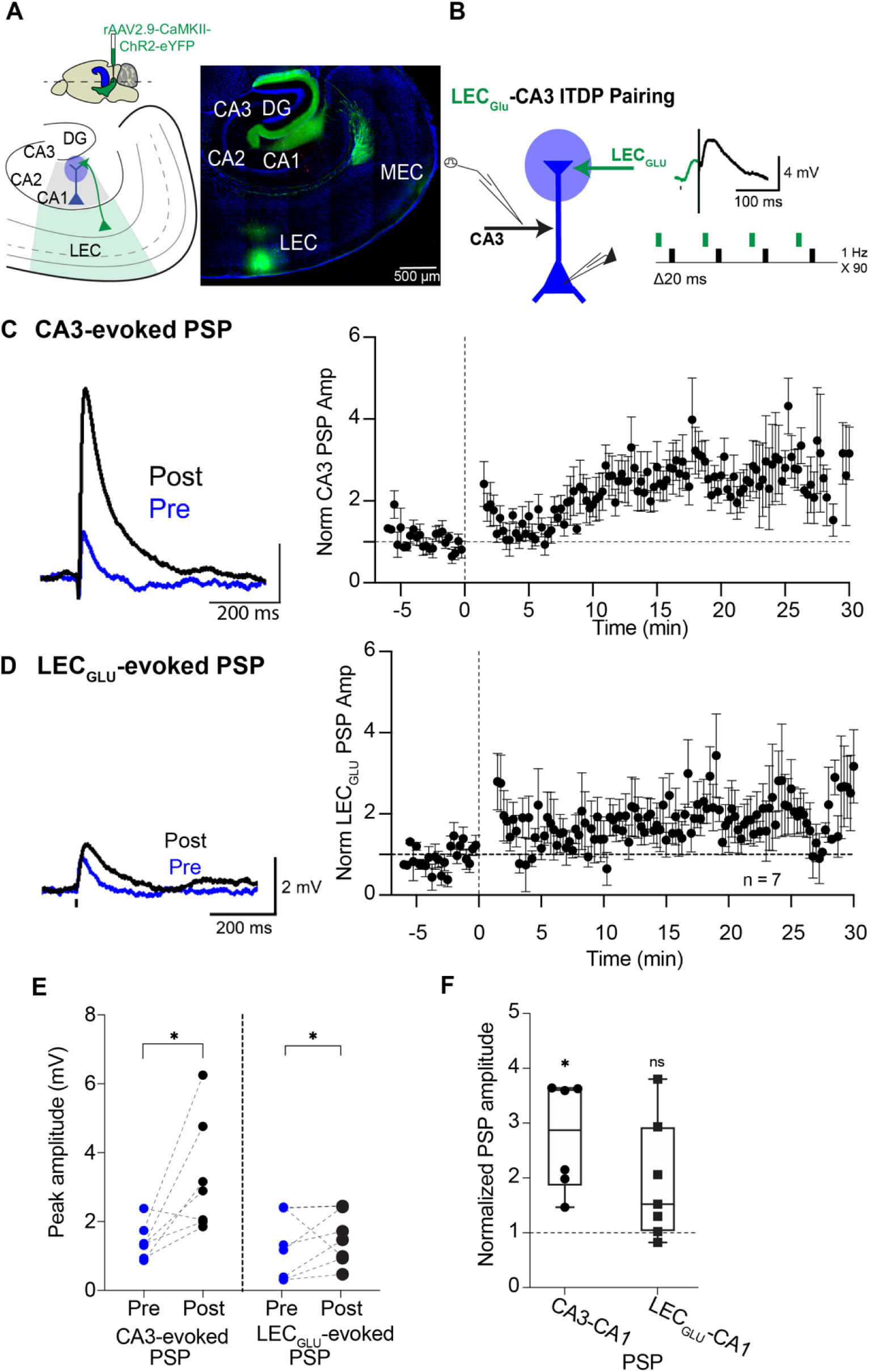
Expression of Input-Timing Dependent Plasticity (ITDP) in area CA1 requires activation of LEC glutamatergic inputs and CA3 Schaffer Collaterals. **(A)**. Left, Schematic representation of the injection strategy for optogenetic activation in LEC (top) and schematic of slice preparation with LEC and CA1 (bottom). Right, Confocal image of the injection site in LEC and the LEC_Glu_ projections in hippocampus. **(B).** CA1 circuit diagram of experimental design. Optogenetic stimulation at the LEC_GLU_ inputs (LEC_GLU_, blue circle and blue arrow) and stimulating electrode at Schaffer collateral from CA3 (CA3, black arrow) excite distal and proximal dendrites of CA1 pyramidal neuron. Right, opto-electrical stimuli design to induce ITDP with optogenetic stimuli of LEC_GLU_ and CA3 electrical stimuli with 20ms interval at 1Hz 90s (see trace example).. **(C).** Time course of ITDP from CA3 PSP (mean ± SEM). ITDP was measured by normalizing the averaged SC PSP measured 30 min after induction of ITDP from the baseline PSP value before pairing. Traces show normalized CA3 PSP pre-(blue) and post-(black) ITDP induction. Dashed lines from time course correspond for x axis time point of stimulation and y normalization of 1, all data points above 1 are measured as ITDP. **(D).** Time course of ITDP of the LEC glutamatergic PSP (mean ± SEM). ITDP was measured by normalizing the averaged CA3 PSP measured 30 min after induction of ITDP from the baseline PSP value before pairing. Traces show normalized CA3 PSP pre-(black) and post-(blue) ITDP induction. Dashed lines from time course correspond for x axis time point of stimulation and y normalization of 1, all data points above 1 are measured as ITDP. Mean potentiation is average of the last 10 minutes post-induction (2.923 ± 0.3063 SEM for CA3-CA1; 1.776 ± 0.04743 SEM for LEC-CA1).. **(E).** Role of LEC glutamatergic inputs in ITDP induction and expression. Individual experiments (circles) measured as ±SEM of normalized CA3 and LEC_GLU_ PSPs before (pre, blue circles) and after (post, black circles) induction of ITDP. Each data point represents one experiment. Statistical significance is a paired two-tailed Wilcoxon test with a significance in -pre and -post for CA3 evoked response (*p=0.0312), and -pre - post for LEC_GLU_ evoked response (*p=0.0469).. **(F).** Plot of individual and min to max of normalized PSP amplitude of CA3 PSP and LEC glutamatergic IPSP. Each group was tested for variation from the theoretical mean of 1 (chance value) using a two-tailed one-sample t-test. For CA3-CA1 **p=0.0075, for LEC_GLU_-CA1 ns p=0.0665. n=7. Data are present as mean ± SEM.

## Discussion

Interactions between LEC and hippocampus have been linked to processing of non-spatial, behavioral relevant information, including association of sensory cues with reward or aversion and temporal sequences ^42^. However, the contribution of LEC input into hippocampal function during different phases of memory, particularly during recall, remains poorly understood^10–13,42^.

Here, we show that LEC glutamatergic projections (LEC_GLU_) to dorsal CA1 are preferentially engaged during memory recall in spatial navigation and object recognition-based tasks. Fiber photometry recordings of LEC_GLU_ axons in CA1 revealed robust and task-relevant activity during recall, indicating an active role for these inputs beyond initial learning. Consequently, pathway-specific chemogenetic silencing of LEC_GLU_ projections in CA1 impaired recall performance, demonstrating that this circuit is necessary for the stable expression of learned information. To further investigate the underlying synaptic mechanisms, performing ex-vivo electrophysiological recordings in CA1. We found that coincident, precisely timed activation of LEC_GLU_ inputs and CA3-Schaffer collaterals are required to induce a non-Hebbian heterosynaptic mechanisms of ITDP. Previous studies have described robust presynaptic short-term facilitation of the perforant path inputs, particularly to CA1 pyramidal neurons^43^. Here we show that LEC_GLU_ is not only effective in inducing LTP in the intra-hippocampal CA3-CA1 pathway but also undergoes lasting homosynaptic potentiation. Together, these findings identify the LEC_GLU_ projections in CA1 as a key component of the circuit mechanisms supporting memory retrieval and challenge the view that LEC function is restricted to non-spatial processing. This supports the idea that there is more functional crosstalk between the MEC and LEC – canonically the “where” and “what” streams of information than previously believed.

MEC and LEC were considered two anatomically, molecularly, and functionally distinct entities of entorhinal cortex^44–46^. MEC undoubtedly plays a role in spatial coding, as evidenced by the presence of grid,^47,48^ head-direction,^49^ and border^50,51^ cells functioning as the “where” of events^52^. In conjunction, LEC shows weak spatial selectivity but has context-dependent responses^53^ to odor,^12,54^ timing^55^, task sequence^42^, novelty,^53^ and objects and their displacement, working as the “what” of experiences.^11^ Counterintuitively, partial and complete MEC lesions and genetically targeted have only mild impacts on hippocampal place coding^21,56^ and appear to be ineffective at altering behavioral performance in some types of spatial navigation tasks^56–58^ and rather have a pronounced effect in temporal associative memory^1,52,59^. Surprisingly, LEC lesions do result in a deficit in place cell rate remapping in area CA3, though their basic spatial firing properties were unaltered^60^. Cross interactions between the spatial and contextual information processing systems are likely mediated by direct glutamatergic and GABAergic projections from LEC to MEC into layer 1 and vice versa^61,62^, as well as feedback from the hippocampus to EC^63^, which carries mixed information from LEC and MEC integrated within the hippocampus^46,64,65^.

At the circuit level, LEC sends both glutamatergic and GABAergic long-range projections to the hippocampus which engage distinct microcircuits across subregions^25,30,41,66^ and modulate distinct sub-compartment specific and functional outcomes. In CA1, LEC excitatory input engages VIP+ disinhibitory neurons that suppress dendrite-targeting inhibitory neurons (INs), including CCK+ and SST+ INs while the GABAergic input directly disinhibits dendrite targeting INs as well^25,41^. Effectively, the LEC glutamatergic and GABAergic inputs to CA1 facilitate dendritic spike generation, and induction of plasticity at the cellular and circuit levels. Functionally this manifests as a gate for encoding contextually salient information, and as shown by our current study, supports retention and recall of long-term memory about both objects and place. On the other hand, in CA3, LEC glutamatergic and GABAergic input conjunctively modulate recurrent collateral driven somatic excitability via distinct soma-targeting CCK+ and VIP+ basket cells^25,30,41^. Functionally, in CA3, silencing the LEC glutamatergic and GABAergic inputs both impact learning but not recall in a head-fixed goal directed spatial navigation task. These pathway and subregion-specific mechanisms may underlie the differential contributions of LEC input to learning versus recall. In this framework, our findings support a model in which LEC to CA1 glutamatergic projections are preferentially recruited during recall to modulate hippocampal output and stabilize memory representations once learning has occurred. What mechanisms could underlie this recall finding? In CA1, temporally precise pairing of entorhinal and Schaffer collateral inputs can drive dendritic spikes and NMDA/mGluR-gated input timing dependent plasticity, supporting both rapid and long-lasting synaptic changes and phosphorylation and kinase dependent transcriptional and translational cascades, underlying early and late phases of LTP^25,41,67^. The activity observed here may reflect recruitment of these mechanisms to reinforce hippocampal ensemble connectivity by synapse strengthening to support high fidelity reactivation of context dependent ensembles during recall. While neuromodulatory inputs especially dopamine and norepinephrine, which affect short-term and long-term plasticity of LEC input to hippocampus ^68,69^ and heavily project to and influence LEC activity during complex memory association behaviors^70^ may regulate these processes. Their precise contribution in this context remains to be determined.

Given the critical role of LEC to CA1 glutamatergic projections in recall, disruption of this pathway may contribute to memory deficits in disease^71–76^. The LEC is among the earliest regions affected in Alzheimer’s disease, and impaired entorhinal input may compromise hippocampal ensemble stability during recall^71–73,75^. These findings provide a circuit level framework for understanding early memory dysfunction across neurological conditions.

## Methods

### Animals

All experiments were conducted in accordance with the National Institutes of Health (NIH) guidelines and with the approval of the New York University Grossman School of Medicine Institutional Animal Care and Use Committee (IACUC). Mice were obtained from The Jackson Laboratory (JAX), and subsequent breeding was established in-house. For anatomical tracing, freely moving fiber-photometry experiments and chemogenetic silence experiments, CAMKII-Cre (JAX, 005359) mice were used to label excitatory neurons or to maintain the same background among all experiments. Mice of both sexes, >12 weeks old up to 26 weeks were used and were housed maintained under a 12:12-h light/dark inverted cycle, with lights turned off at 10:00 am and turned on at 10:00 pm in a controlled temperature at 22 ± 2°C with *ad libitum* access to food and water.

### Viruses

For anatomical tracing of axonal density from LEC into dorsal hippocampal area CA1 experiments, we unilaterally injected AAV1-Syn-Flex-NES-jRGECO1a-WPRE-SV40 (a gift from Douglas Kim & GENIE Project; Addgene viral prep # 100853-AAV1; http://n2t.net/addgene:100853; RRID: Addgene_100853) in 6 mice into LEC of CaMKII-Cre mice. To verify that the density of projections is independent of the viral expression, we injected in a subset of 2 mice AAV5-CaMKII-Chronos-GFP (Neurophotonics no. 319) with no differences observed. For in vivo fiber photometry recordings in freely moving experiments, we injected AAV1-Syn-Flex-NES-jRGECO1a-WPRE-SV40 (a gift from Douglas Kim & GENIE Project; Addgene viral prep # 100853-AAV1; http://n2t.net/addgene:100853; RRID: Addgene_100853) unilaterally into LEC of CAMKII-Cre mice. The RGeCO1a virus was incubated for at least 21 days when we observed functional expression in our fiber photometry system. Behavioral tasks were started once the signal was observed and performed during 12 days after the first indication of calcium signal. For in vivo chemogenetic silencing in freely moving cognitive behavioral experiments, we injected AAV5-CAMKII-PSAM4-GlyR-IRES-EGFP (was a gift from Scott Sternson; Addgene viral prep # 119744-AAV5; http://n2t.net/addgene:119744; RRID: Addgene_119744) bilaterally into LEC of CaMKII-Cre mice. The PSAM4 virus was incubated for at least 37 days (5 weeks) to verify axonal expression in hippocampal area CA1. Freely moving behavioral tasks were started after the 5-week period and performed during 15 days after. In a subset group of mice for analysis of control conditions, we injected AAV5-CaMKIIa-EGFP (a gift from Bryan Roth; Addgene viral prep # 50469-AAV5; http://n2t.net/addgene:50469; RRID: Addgene_50469) bilaterally into LEC of CaMKII-Cre mice. The EGFP virus was incubated for at least 37 days (5 weeks) to verify axonal expression in hippocampal area CA, like the PSAM viral injection. For ex-vivo experiments, AAV2.9-CaMKIIa-hChR2(H143R)-eYFP was injected into LEC (a gift from Karl Deisseroth; Addgene viral prep #26969-AAV9; http://n2t.net/addgene:26969; RRID: Addgene_26969) unilaterally into LEC of CaMKII-Cre mice. The CHR2 virus was incubated for 7 to 10 days.

### Stereotaxic surgeries

*To express viral constructs, Chronos, RGeCO1a, PSAM4, EGFP and CHR2 into LEC glutamatergic neurons.* Mice were put under anesthesia with isoflurane (1 to 2.5%, inhalation) and injected with buprenorphine (0.1 mg/kg, intraperitoneal IP, Covertus). After, mice were placed and head fixed in a stereotaxic apparatus (Stoeling), eye ointment was applied and exposition of head skin was prepared with previous shaving and disinfection with ethanol (at 70%) and betadine. We applied ropivacaine hydrochloride topically (1 to 2 drops, 2 mg/ml; Hospira). A small incision was made to expose the skull and hydrogen peroxide (0.1%) was applied to clean and increase the visualization of cranial sutures. Once skull was fixed, viral injection pipettes were filled with mineral oil and with the virus using a Nanoject II injector (Drummond Scientific). The skull was leveled according to bregma and lambda alignment along the z axis. A craniotomy was drilled in the skull with a dental drill (Foredom Electric) and verification of z-coordinate without the pia damage or interference was zeroed on the flat surface. 276 nl of virus was injected into the brain at coordinates as it mentions in table 1. The pipette was slowly lowered into the tissue up to 0.1 mm deeper than the deepest coordinate, left at this location for 30s, retracted to the first coordinate and held there for 1 min prior to injection. 46 nl of virus was injected in two increments of 23 nl spaced by 15 s at each z coordinate, with an additional 3 to 5 min pause between z coordinates and a final 10 min wait at the deepest (last) z coordinate of each injection site before the retraction of the injection pipette outside of brain tissue. Once all coordinates were properly injected, mice were sutured, giving Neosporin topically on the surgical incision, rehydrate with 1 ml of sterile saline subcutaneously and monitored until full recovery from anesthesia was observed. For postoperative care, intraperitoneally injection of buprenorphine 0.1 mg/kg for 3 days were injected. Infection sites were confirmed as a post-hoc after all the behavioral tasks were evaluated.

**Table 1.** Coordinates for viral injection in LEC.

| **Lateral Entorhinal Cortex (LEC)** |  |  |
| *Axis of injection* | *Coordinates of Injection* |  |
|---|---|---|
| Medial-lateral relative to midline (x) | ± 4.5 | ± 4.7 |
| Anterior-posterior relative to bregma (y) | -3.2 | -3.0 |
| Dorsal–ventral relative to surface of the brain (z) | -2.5 | -2.5 |
|  | -2.8 | -2.8 |
|  | -3.0 | -3.0 |

### For optical fiber cannulae implants in dorsal CA1

To record the glutamatergic axons from LEC into dorsal CA1 during freely moving fiber photometry recordings, we obtain raw calcium transients signals through a 400 μm optic fiber (Thorlabs, FT400EMT) coupled to a ceramic ferrule (Thorlabs, CFLC440) implanted 50 μm above SLM in dorsal CA1. The optical fibers were polished by sanding the tip for optimal light transmission, using fiber polishing film of a decrease grit size (Thorlabs, LF30D (30 μm) > LF6D (6 μm) > LF3D (3 μm) > LF1D (1 μm) > LFCF (0.02 μm). Fiber tip quality was checked with a fiber inspection scope (Thorlabs, FS201). Optic fibers only with a clean tip surface that transmitted at least 80% of intensity of LED (only 20% of leak was permitted, using an optical power meter; Thorlabs, PM100D) were used. Surgeries were done 2 weeks after viral injection and the procedures to prepare mice skull were identical to that of the stereotaxic viral injections. Once skull stayed fixed, we used a straight clamp with pad (Kopf, 1271-C-MOD) to hold the optical fiber assembly onto the electrode holder used during the virus injection. We drilled a hole and lowered the fiber optic implant into the brain unilaterally into the same viral injection hemisphere at the following coordinates for dorsal CA1from table 2. Once secured into the coordinates the implant was secured to the mouse brain using dental cement (Parek). After implantations, animals were injected with 1.0 ml saline solution to rehydrate. Mice were given intraperitoneally injections of 0.1 mg/kg of buprenorphine for 3 consecutive days following surgery recovery.

**Table 2.** Fiber photometry Cannulae placement.

| **Dorsal CA1** |  |
| *Axis of injection* | *Coordinates of Injection* |
|---|---|
| Medial-lateral relative to midline (x) | ± 1.5 |
| Anterior-posterior relative to bregma (y) | -2.2 |
| Dorsal–ventral relative to surface of the brain (z) | -1.65 |

### For dual-bilateral implants of cannula guide in dorsal CA1

To silence selectively the glutamatergic projections from LEC into dorsal CA1, we used local infusion of the cognate synthetic ligand PSEM817 in CA1 using a bilateral cannula as previously reported ^25^. The surgical and craniotomy procedures were identical as of the stereotaxic viral injections. A sterilized custom designed bilateral cannula guide was inserted in the skull over dorsal hippocampal area CA1as follows in table 3 along with a dummy cannula and cover with a custom-made dust cap. The implant was secured to the skull with dental cement (C & B Metabond) and all the surrounding skull area to secure the implant. The cement was allowed to dry for at least 20 min or when verification of dryness was observed.

**Table 3.** Dual-bilateral implants cannulae placement.

| **Dorsal CA1** |  |
| *Axis of injection* | *Coordinates of Injection* |
|---|---|
| Medial-lateral relative to midline (x) | ± 1.5 |
| Anterior-posterior relative to bregma (y) | -2.2 |
| Dorsal–ventral relative to surface of the brain (z) | -1.5 |

### Immunohistochemistry

As previously reported^77^, once in vivo experiments were finished, mice were anesthetized and transcardially perfused with PBS-1X and a solution of paraformaldehyde (PFA) 4% with PBS. Brains were dissected and fixed for 4 hours at 4°C and then sectioned into 400-μm thick coronal slices using a vibratome (Leica VT1000S). Slices went into tissue clearing technique CUBIC^78,79^, they were washed with 0.1 M PB and then immersed in CUBIC-1 solution for 2 days. Next, slices were washed in 0.1 M PB and went into standard immunostaining protocol, the slices were permeabilized in 0.5% PBS-Triton (PBST), blocked in 3% normal goat serum (NGS) overnight and then incubated with primary antibodies overnight at 4°C. Afterwards, slices were washed and incubated with secondary antibodies overnight at 4°C. Once secondary antibodies were incubated, slices were thoroughly washed with 0.1M PB and then incubated with CUBIC-2 solution before being mounted on the same solution. For primary antibodies, we used rabbit polyclonal anti-RFP primary antibody (1:1,000 dilution; Rockland/Thermo Fisher, 600-401-379-RTU), chicken anti-GFP (1:1,000 dilution; Abcam, 13970), and rabbit polyclonal anti-GFP primary antibody (1:1,000 dilution; Invitrogen, A-6455). For secondary antibodies, we used goat anti-chicken 488 (1:500 dilution; Invitrogen, A11039), goat anti-rabbit 488 (1:500 dilution; Invitrogen, A11008), goat anti-rabbit 555 (1:500 dilution; Invitrogen, A21428) and neurotrace 435/455 (1:200 dilution; Invitrogen, N21479). Once mounted, slices were imaged using an inverted Zeiss Axio Oberserver Z.1 confocal microscope, using (magnification of 10X air immersion) and 20X air-immersion objectives (Zeiss) and 405-nm, 488-nm, and 594-nm lasers to acquire 16-bit image stacks. Stacks were acquired every 5 μm, tiled images were stitched using Zen 2.6 Blue Microscopy Software (Zeiss), and the tiled z-stacks were subsequently processed using Fiji (ImageJ).

### Fiber Photometry Recordings

To record RGECO1a signals, we used a fiber photometry system based on a Doric Lenses photometry console (Doric Lenses, Québec, Canada) controlled by Doric Neuroscience Studio software (version v6.3.2.0). Excitation light was sinusoidally modulated, and fluorescence sampled at 60.236 Hz. Excitation and emission separation were achieved with a fluorescence minicube (Doric Lenses, model FMC6; configured for 555–570 nm emission and a 405 nm isosbestic channel), which routed excitation light to and collected emitted light from the implanted optical fiber. Excitation for the calcium dependent RGECO1a signal was provided by a 565 nm fiber coupled LED (M565F3, Thorlabs), and a 405 nm fiber coupled LED (M405F3, Thorlabs) was used as an isosbestic control to monitor movement related artifacts. Both LEDs were driven and modulated by the Doric console via an LED driver (Thorlabs, model LEDD1B); the 565 and 405 nm channels were sinusoidally modulated at distinct frequencies and demodulated in real time by the console. LEDs were coupled to the minicube using low autofluorescence multimode patch cords (Doric Lenses; 400 µm core, 0.40 NA, FC/PC connectors). The combined excitation light was delivered to the brain through a similar patch cord connected via a rotary joint (Thorlabs, model RJ1 with 400 - 700 nm Wavelength Range) to the implanted fiber optic cannula (Thorlabs; 400 µm core, 0.40 NA). Excitation power at the tip of the patch cord/implanted fiber was adjusted to minimize photobleaching and tissue heating. Fluorescence emitted by RGECO1a was collected through the same fiber, returned to the minicube, spectrally separated by dichroic mirrors and band pass filters (centered at 555–570 nm), and directed to a Newport Femtowatt silicon photoreceiver (Newport; 320–1050 nm). The analog voltage output from the photoreceiver was digitized by the Doric console and stored for offline analysis. The 405 nm isosbestic channel was acquired in parallel and visually inspected to identify movement related artifacts but was not used for ΔF/F correction.

For preprocessing, raw fluorescence traces were low-pass filtered (2nd-order Butterworth) and detrended to correct for slow photobleaching using a single-exponential fit. Baseline fluorescence (F₀) was estimated using a sliding window approach, and relative fluorescence changes were computed as ΔF/F = (F – F₀)/F₀. Trials or epochs containing saturated signals, abrupt step changes, or large motion artifacts (identified from the raw and 405 nm signals) were excluded from further analysis. Behavioral video was acquired at 24 fps using a Basler camera (model acA1300-60gmNIR), and TTL pulses from the Doric console were used to hardware-synchronize video and photometry recordings for precise temporal alignment. All preprocessing and analyses were performed using custom-written scripts in MATLAB and Python. For preprocessing, raw fluorescence traces were low-pass filtered (2nd-order Butterworth) and detrended to correct for slow photobleaching using a single-exponential fit. Baseline fluorescence (F₀) was estimated using a sliding window approach, and relative fluorescence changes were computed as ΔF/F = (F – F₀)/F₀. Trials or epochs containing saturated signals, abrupt step changes, or large motion artifacts (identified from the raw and 405 nm signals) were excluded from further analysis.

### In vivo Behavior

Like previously reported^77^. Mice were tested in two behavioral paradigms, novel object recognition (NOR) and the Barnes maze (BM). Axonal activity was recorded using fiber photometry through the whole behavioral paradigms, while chemogenetic silencing was restricted to recall phases of each task.

Behavior was recorded with a Basler camera (acA640-100gmNIR, 24fps) coupled to pylon Camera Software, and exploration was scored offline as time spent actively investigating each object (head and snout oriented toward and within ∼1–2 cm of the object). For photometry recordings, behavioral video recordings were processed with DeepLabCut^80^ to track position in XY of animals’ head to then aligned with the photometry recordings for post-processing. Photometry and video recordings were hardware synchronized via TTL pulses from the Doric console to allow precise alignment of neural activity with object exploration bouts during encoding and recall. Analysis for the photometry signal was performed through a custom made MATLAB script and then synchronized with the behavioral features with a Python custom made scripts. For chemogenetic silencing, behavioral analysis was performed and tracked using ANY-maze video tracking software program (ANY-maze Stoeling, 4.99m application version, 4.115 data file version).

#### Novel Object Recognition (NOR)

NOR was performed in an open-field arena (length 37 cm, width 29 cm, height 27 cm) with opaque walls under dim illumination. On the test day, mice were placed in the arena containing two identical objects and allowed to freely explore for 10 min (encoding session) while calcium signals were recorded via fiber photometry or without any chemogenetic strategy. Animals were then returned to their home cage for a 30 min delay. During the subsequent recall session, one of the familiar objects was replaced with a novel object, and mice were allowed to explore for another 10 min with concurrent photometry recording. For chemogenetic silencing experiments with the PSAM-PSEM817 system, the designer drug (PSEM817; 1 mg/kg,15 µM intracerebral) or vehicle was administered 5 min prior to the recall session only; the encoding session was always conducted drug□free. Object identity (familiar vs novel) and spatial position (left vs right) were counterbalanced across animals. For analysis, we evaluated the time of exploration of head and snout oriented, and the indices were evaluated as follows:

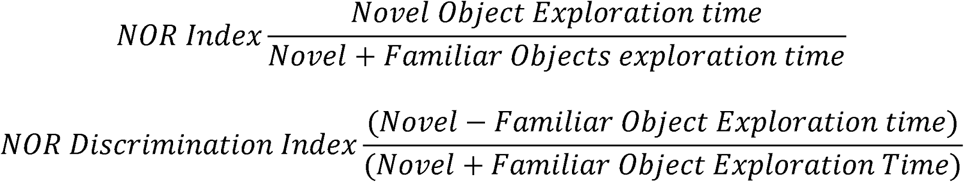

#### Barnes maze (BM)

Spatial learning and memory were assessed in a circular Barnes maze (90 cm in diameter) with 20 equally spaced holes around the perimeter, one of which was connected to an escape box ^81^. Distal visual cues were placed around the room and kept constant across days. Mice underwent 7 consecutive learning days, with 4 trials per day (maximum trial duration: 3 min; inter trial interval: 15 min). The escape hole location remained fixed for each animal. Fiber photometry was recorded in all trials. Fiber photometry analysis, learning and recall were grouped into four stages (see Fig. 3b): Stage 1 (early learning: days 1–2, trials 1–2), Stage 2 (days 3–4, trials 1–2), Stage 3 (late learning: days 6–7, trials 1–2), and Stage 4 (recall: day 7, trial 3-4 and day 8, trial 1). On day 8, mice performed a single recall trial with the escape box in the same location as during training. For chemogenetic silencing experiments, the designer drug PSEM817 (15 µM) or vehicle (saline 0.9%) was administered 5 min prior to day 8 recall trial only; all learning trials were conducted drug□free.

During each Barnes maze trial, mice were placed in the center of the maze and allowed to explore until they entered the escape box or the trial ended. Behavior was recorded with the same Basler camera setup used for NOR. Hole nose pokes and escape hole interactions were scored offline, and latency to locate the escape hole, path trajectories, and search strategies were quantified. Video timestamps and TTL synchronization pulses from the Doric console were used to precisely align behavioral events (e.g., nose pokes at escape vs non escape holes) with calcium signals across the four stages of learning and recall. For chemogenetic silencing, analysis was evaluating the number of errors, representing the number of times the mice explored holes including the escape hole without entering and latency represents the total time that the animal takes to enter the escape hole.

### Slice preparation and Electrophysiology

As our previous studies^38,41^ thick (400 μm) horizontal hippocampal slices of brains from mice that were transcardially perfused with ice-cold sucrose-enriched dissection artificial cerebrospinal fluid (dACSF) were obtained with a vibrating microtome (Leica VT1200S). The slices were incubated in a 1:1 mixture of sucrose-enriched dACSF and standard ACSF solution for 15 min at 34°C and 45 min at room temperature before being transferred to the recording chamber containing standard ACSF. Infrared- or fluorescence-guided whole-cell patch-clamp recordings were performed at 34°C in standard ACSF. Fire-polished borosilicate glass pipettes (Sutter) were used with tip resistances of 3.5–4.5 MΩ for somatic recordings. A Multiclamp 700B Amplifier coupled to a Digidata digitizer and pClamp 9 software (Axon Instruments) were used for data acquisition. The average series resistance for whole-cell voltage-clamp recordings was kept between 9–15MΩ; 75%–80% of the resistance was compensated. Current-clamp recordings in KMeSO4 intracellular recording solution were obtained with access resistances of 10–20 MΩ for the soma, compensated in bridge mode.

To measure post-synaptic potential (PSP) responses, CA1 neurons were kept at their resting membrane potential (RMP). Optogenetic stimulation was performed using a Thorlabs LED system (405/470/565/635 nm wavelengths). ChR2^+^ LEC Glutamatergic (LEC_GLU_) inputs were activated using a single 2 ms light pulse (470 nm, every 15 s) delivered at <50% maximum photostimulation strength (23 mW, 50 µm diameter beam spot). The CA3 inputs were electrically stimulated using focal glass pipette-stimulating electrode placed in stratum radiatum of CA1 coupled to constant current stimulator (WPI). The stimulation strength for LEC_GLU_ light-evoked responses and CA3 electrically evoked were adjusted to be roughly <50% of maximum amplitude at baseline (typically <2 mV LEC_GLU_ PSP, <4 mV for CA3). Light-evoked responses from LEC and electrically stimulated responses from CA3 Schaffer Collaterals were recorded every 15 s, for baseline recordings (LEC before CA3, 2 s apart) before and after ITDP induction. ITDP was induced by pairing photostimulation of LEC_GLU_ inputs and electrical stimulation of CA3 inputs at a 20 ms interval (LEC_GLU_ before SC, 20 ms apart) at 1 Hz for 90 s.

#### Solutions

Sucrose-enriched dissection ACSF (dACSF) consisted of (in mM): Sucrose 195.0, NaCl 10.0, Glucose 10.0, NaHCO_3_ 25.0, KCl 2.5, NaH_2_PO_4_ 1.25, Sodium pyruvate 2.0, CaCl_2_ 0.5, MgCl_2_ 7.0, saturated with 95% O_2_ and 5% CO_2_. Standard artificial cerebrospinal fluid (ACSF) consisted of (in mM): NaCl 125.0, Glucose 22.5, NaHCO_3_ 25.0, KCl 2.5, NaH_2_PO_4_ 1.25, Sodium pyruvate 3.0, Ascorbic Acid 1.0, CaCl_2_ 2.0, MgCl_2_ 1.0, saturated with 95% O_2_ and 5% CO_2_. The intracellular solution contained the following (in mM): KMeSO_4_ 135 (for current clamp recordings), KCl 5, EGTA 0.1, HEPES 10, NaCl 2, Mg-ATP 5, Na_2_-GTP 0.4, Na_2_Phosphocreatine 10, and Neurobiotin or Biocytin (0.2%). The pH of the intracellular solution was adjusted to 7.2 with KOH as needed. Osmolarity ranged from 270-290 and was adjusted with sterifiltered MilliQ water.

## Extended Figures

**Extended Data Fig 1 (Related to Fig1).**
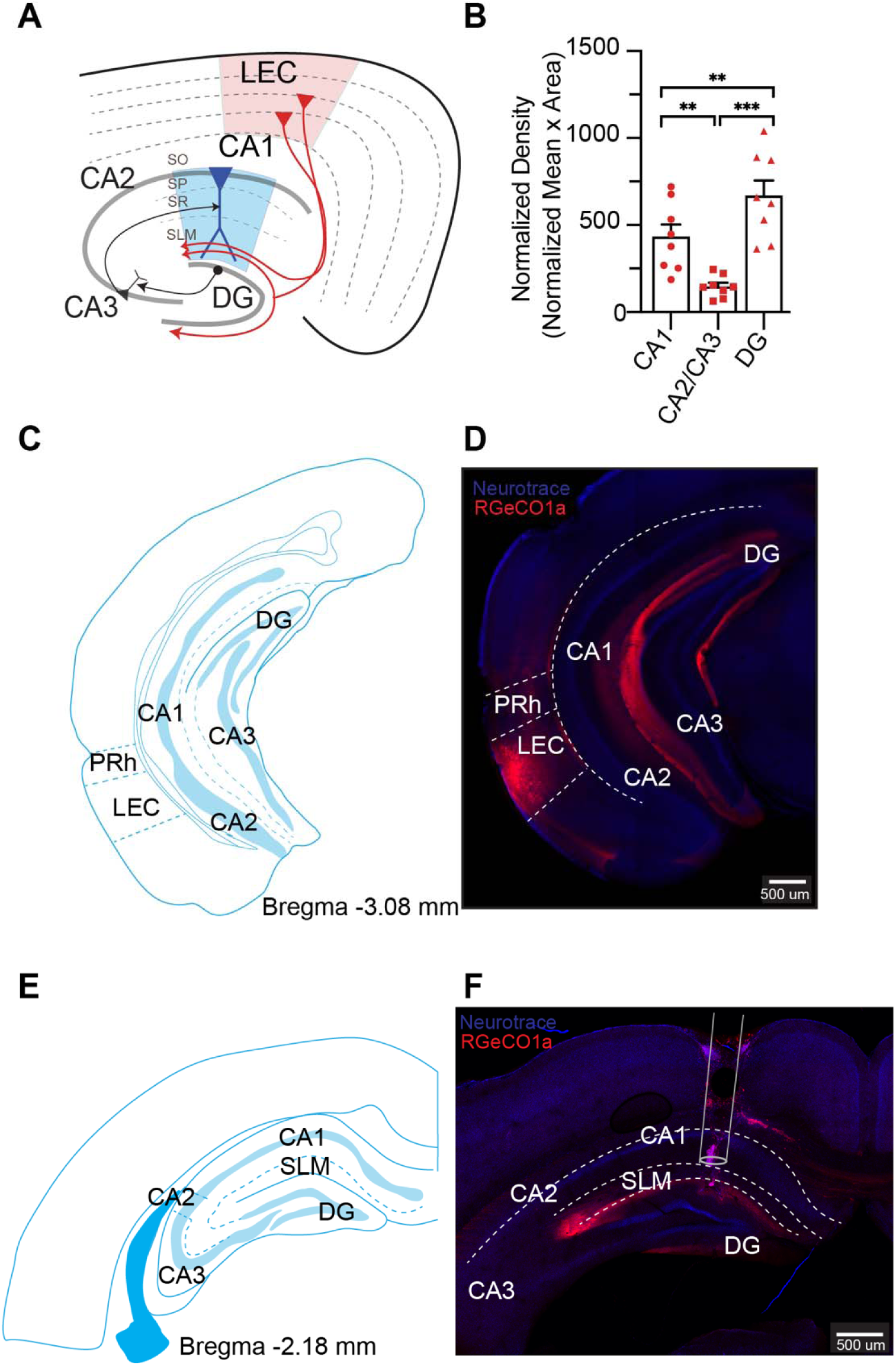
Injection site in LEC with density analysis and cannula placement in area CA1. **(A)** Schematic diagram of horizontal slice showing LEC area (in red) and glutamatergic projections in hippocampal area. **(B)** Quantification of normalized density, analysis of the normalized mean per area from hippocampal area of glutamatergic projections from LEC. A one-way ANOVA was compared to conduct the difference on the density among the hippocampal area (CA1 vs CA2/CA3, **p=0.0025; CA1 vs DG, **p=0.0095; CA2/CA3 vs DG, ***p=0.0005). **(C)** Schematic diagram of coronal section from the brain atlas showing the position of the coronal plane of section through the mouse brain as ventral distance from bregma (−3.08mm) with LEC area. **(D)** Confocal image of coronal brain slice of a mouse injected with AAV-SynFLEX-RGECO1a-mCherry in CaMKII-Cre mice into LEC and immunostained for mCherry (red) and Neurotrace (nuclear marker). LEC glutamatergic neurons are observed and axons in hippocampal area (in red). **(E)** Schematic diagram of coronal section from the brain atlas showing the position of the coronal plane of section through the mouse brain as ventral distance from bregma (−2.18mm) with dorsal hippocampal area. **(F)** Confocal image of coronal brain slice of a mouse with glutamatergic projections from LEC into hippocampal area immunostained from mCherry (red) and Neurotrace (nuclear marker). LEC glutamatergic axons are observed and the track of the cannula above SLM border of area CA1 for fiber photometry recordings. Data are present as mean ± SEM.

**Extended Data Fig 2 (Related to Fig 1).**
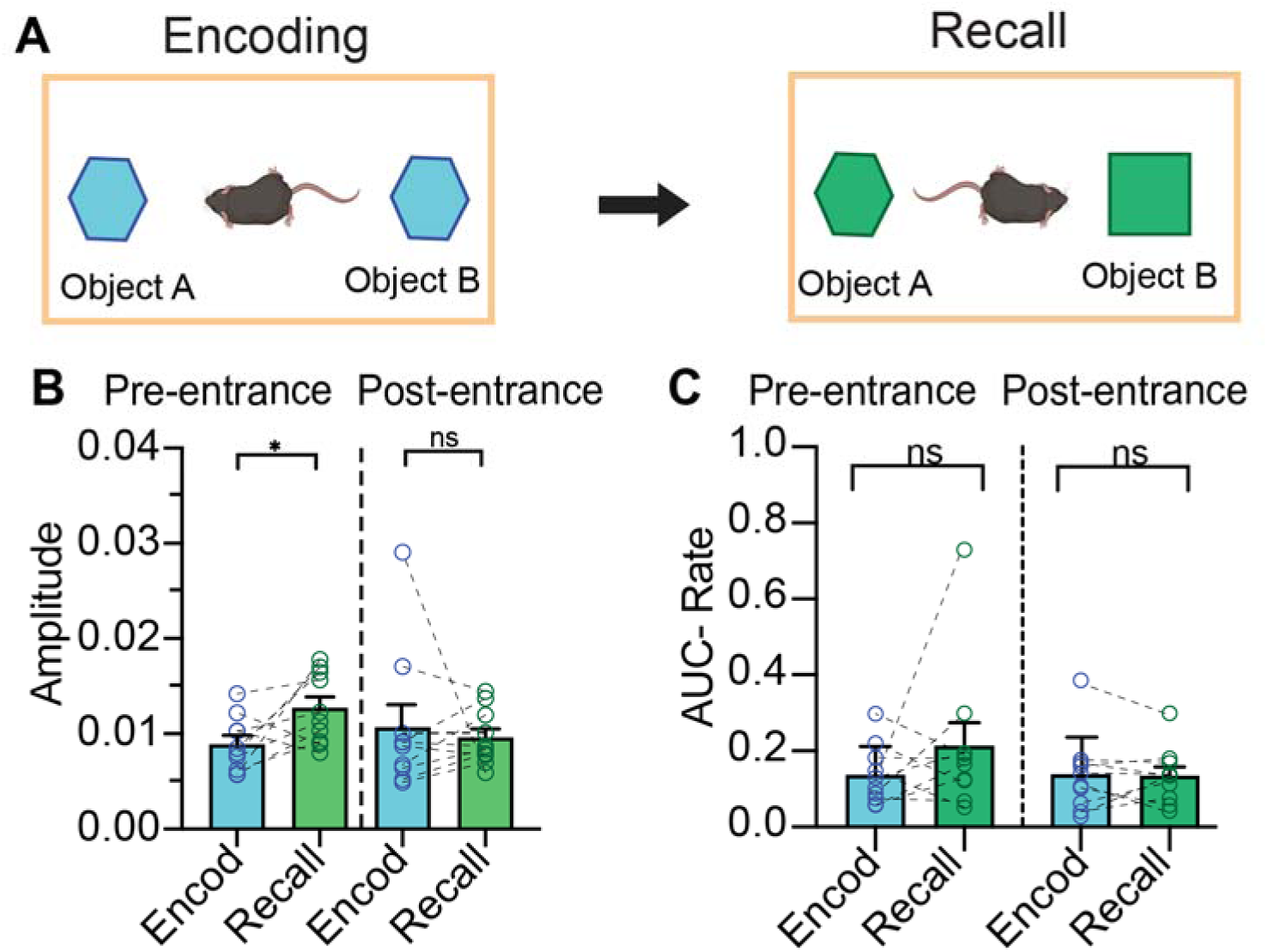
Amplitude and AUC-Rate in encoding and recall. **(A)** Schematic representation of the experimental design for NOR test, first encoding session (blue objects) and recall session (green objects). **(B)** Amplitude of the axonal signal from LEC in CA1 during encoding and recall, 5s before object area entrance and 5s after object area entrance in encoding (blue) and recall (green). Each data point represents one animal and is presented as the SEM. Statistical significance is presented as a paired two-tail t-test with a significance effect between encoding and recall 5s before object area entrance (*p= 0.0339) sessions but not during 5s after object area entrance (p= 0.6923). **(C)** Area Under Curve (AUC) of the axonal signal from LEC in CA1 during encoding and recall, 5s before object area entrance and 5s after object area entrance in encoding (blue) and recall (green). Each data point represents one animal and is presented as the SEM. Statistical analysis is presented as a paired two-tail Wilcoxon test with no significance effect observed 5s before object area entrance (p=>0.9999) and 5s after object area entrance (p=0.8457). Data are present as mean ± SEM.

**Extended Data Fig 3 (Related to Fig 2).**
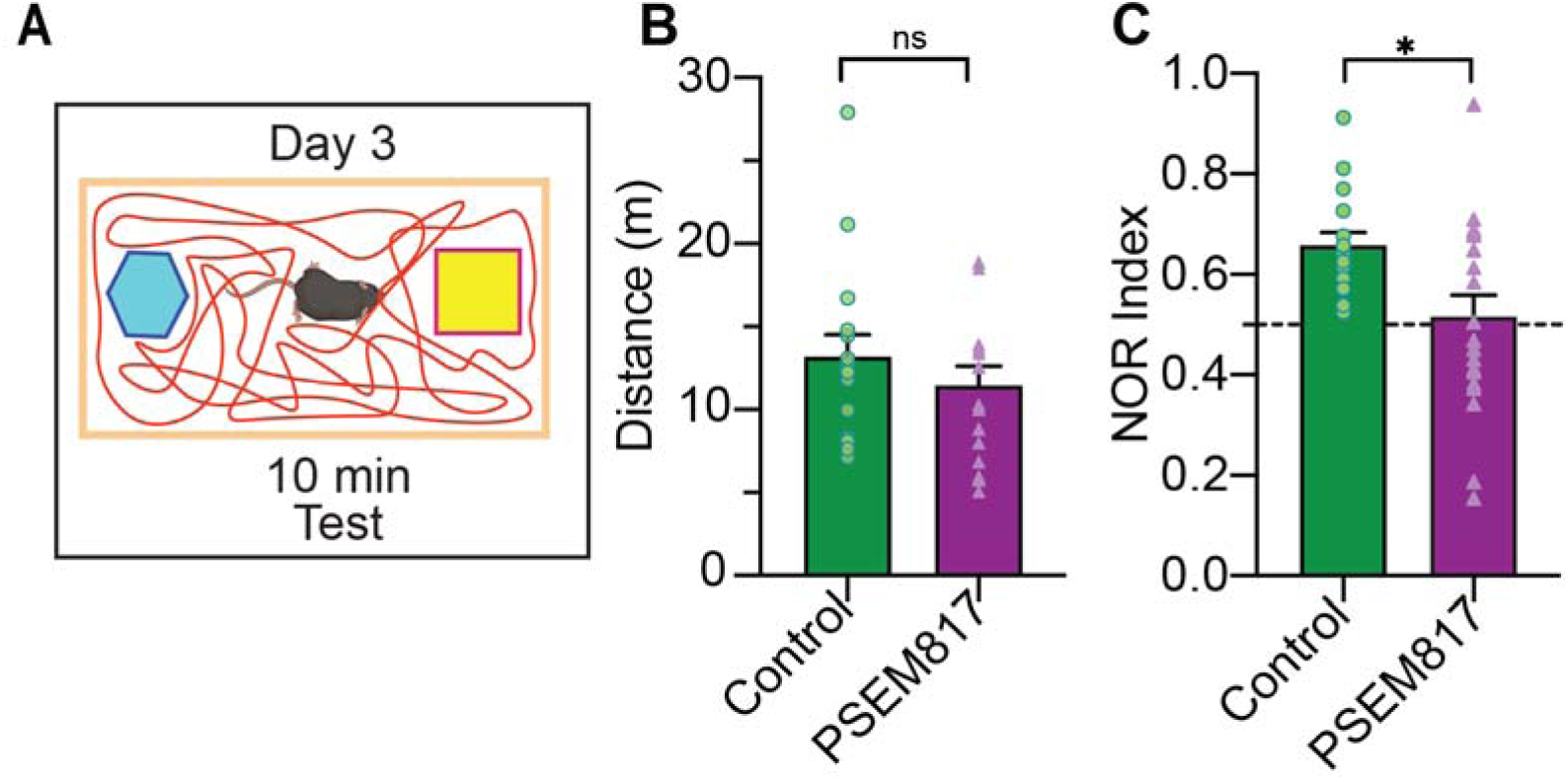
Distance and NOR index during local silencing. **(A)** Schematic representation of the experimental design of NOR recall test showing the familiar object (in blue) and novel object (in yellow) with exemplification of trajectory in red. **(B)** Distance traveled from control and PSEM817 during recall session. Statistical analysis is represented as a SEM in a two-tail U Mann-Whitney test (p=0.4450). **(C)** NOR index from control and PSEM817 during recall session. Statistical analysis is represented as a SEM in a two-tail unpaired t-test (*p=0.0115). Data are present as mean ± SEM.

**Extended Data Fig 4 (Related to Fig 2).**
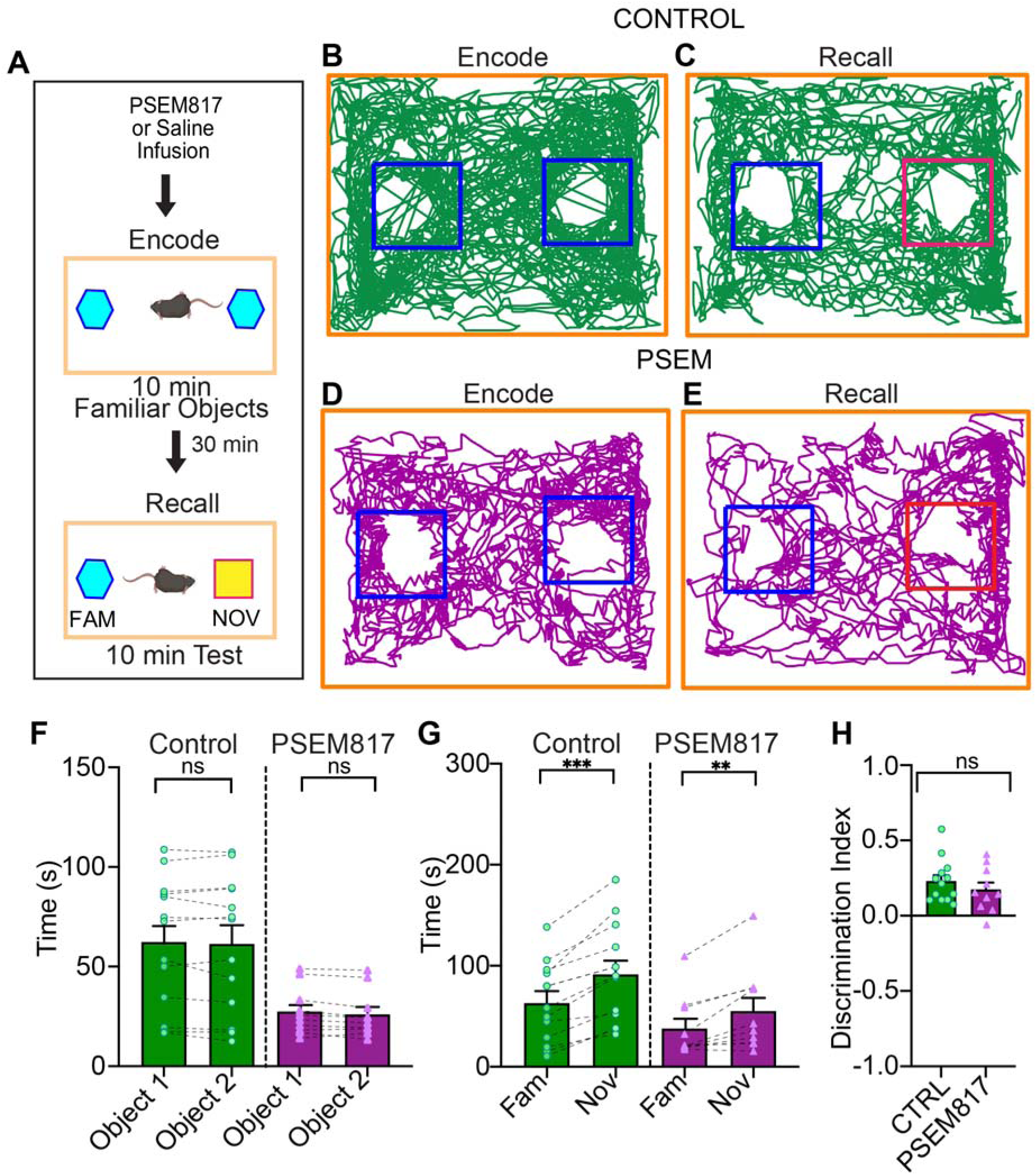
Silencing of glutamatergic LEC to CA1 projections during encoding does not impairs novelty recognition in the novel object recognition test. **(A)** Schematic representation of the experimental design of NOR where silencing is before encoding, showing the familiar object (in blue) and novel object (in yellow. **(B)** Representative map of the head trajectory from one mouse in control conditions (locally infused bilaterally of saline into dorsal area CA1) in encoding of NOR, showing the head distance trajectory and the occupancy (green) into the object area zone from familiar objects (blue square). **(C)** Representative map of the head trajectory from one mouse in control conditions (locally infused bilaterally of saline into dorsal area CA1) in recall of NOR, showing the head distance trajectory and the occupancy (green) into the object area zone from familiar object (blue square) and novel object (red square). **(D)** Representative map of the head trajectory from one mouse in PSEM817 conditions (locally infused bilaterally of PSEM817 into dorsal area CA1), showing the head distance trajectory and the occupancy (dark purple) into the object area zone from familiar objects (blue square). **(E)** Representative map of the head trajectory from one mouse in PSEM817 conditions (locally infused bilaterally of PSEM817 into dorsal area CA1), showing the head distance trajectory and the occupancy (dark purple) into the object area zone from familiar object (blue square) and novel object (red square). **(F)** Analysis of the time spent exploring object 1 and object 2 during encoding trial of NOR in control and PSEM817 with infusion of either saline (control) and PSEM817. For control: 62.29 ± 9.036. object 1, 61.35 ± 9.397 object 2; for PSEM817: 27.51 ± 3.785 object 1, 26.09 ± 3.670 object 2. Statistical significance tested with a two-tailed paired U Mann-Whitney test for both cohorts (control p= >0.9999; PSEM817 p= 0.7394). Analysis of the comparison of time that mice spent exploring the novel and familiar objects in the recall trial of NOR task in control and PSEM817 conditions, no infusion was done. For control; 62.93 ± 10.93 familiar, 91.46 ± 13.43 novel; for PSEM817: 37.85 ± 9.492 familiar, 55.04 ± 12.98 novel. Statistical significance was tested with a two-tailed paired Wilcoxon matched-pairs signed rank test for both controls (control p=0.0002; PSEM817 p=0.0059). **(G)** Comparison of the novel object discrimination index of mice during NOR test trial. Statistical significance tested with a two-tailed unpaired t-test with no changes on the effect of local infusion silencing with PSEM817 during encoding (ns p= 0.3734).

**Extended Data Fig 5 (Related to Fig 3).**
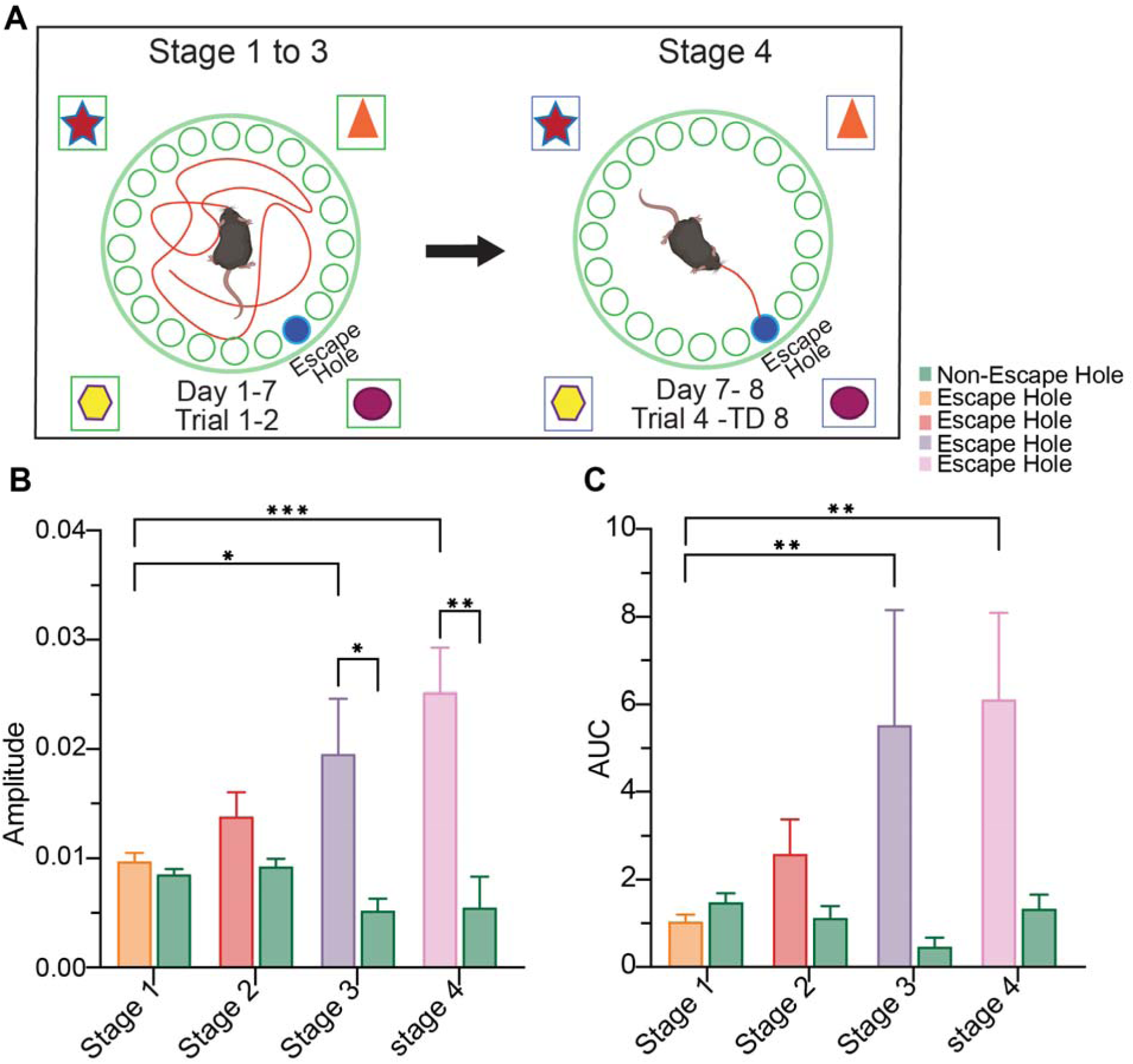
Amplitude and AUC during Stage 1 to Stage 4 related to escape hole area and non-escape holes areas. **(A)** Schematic representation of the experimental design for Barnes maze test, first is the trials from stage 1 to stage 3 (day 1 to day 7, first 2 trials) and next the stage 4 (last trials from day 7 and single trial of day 8) of the test. **(B)** Amplitude of the axonal signal from LEC in CA1 during stage 1 to stage 4 related to escape hole area and non-escape holes areas. Data is presented as SEM and statistical significance is presented as an ordinary Two-Way ANOVA (*p= < 0.05, **p= < 0.01, *** p= < 0.001) with a significant effect between escape hole area in stage 4 and stage 1 and 2 of escape hole area and non-escape hole areas in stage 3 and 4. **(C)** Area Under Curve Rate (AUC) of the axonal signal from LEC in CA1 during stage 1 to stage 4 related to escape hole area and non-escape holes areas. Data is presented as SEM and statistical significance is presented as an ordinary Two-Way ANOVA (*p= < 0.05, **p=< 0.01) with a significant effect between escape hole area in stage 1 and 3 and stage 1 and 4. Data are present as mean ± SEM.

**Extended Data Fig 6. (Related to Fig 4).**
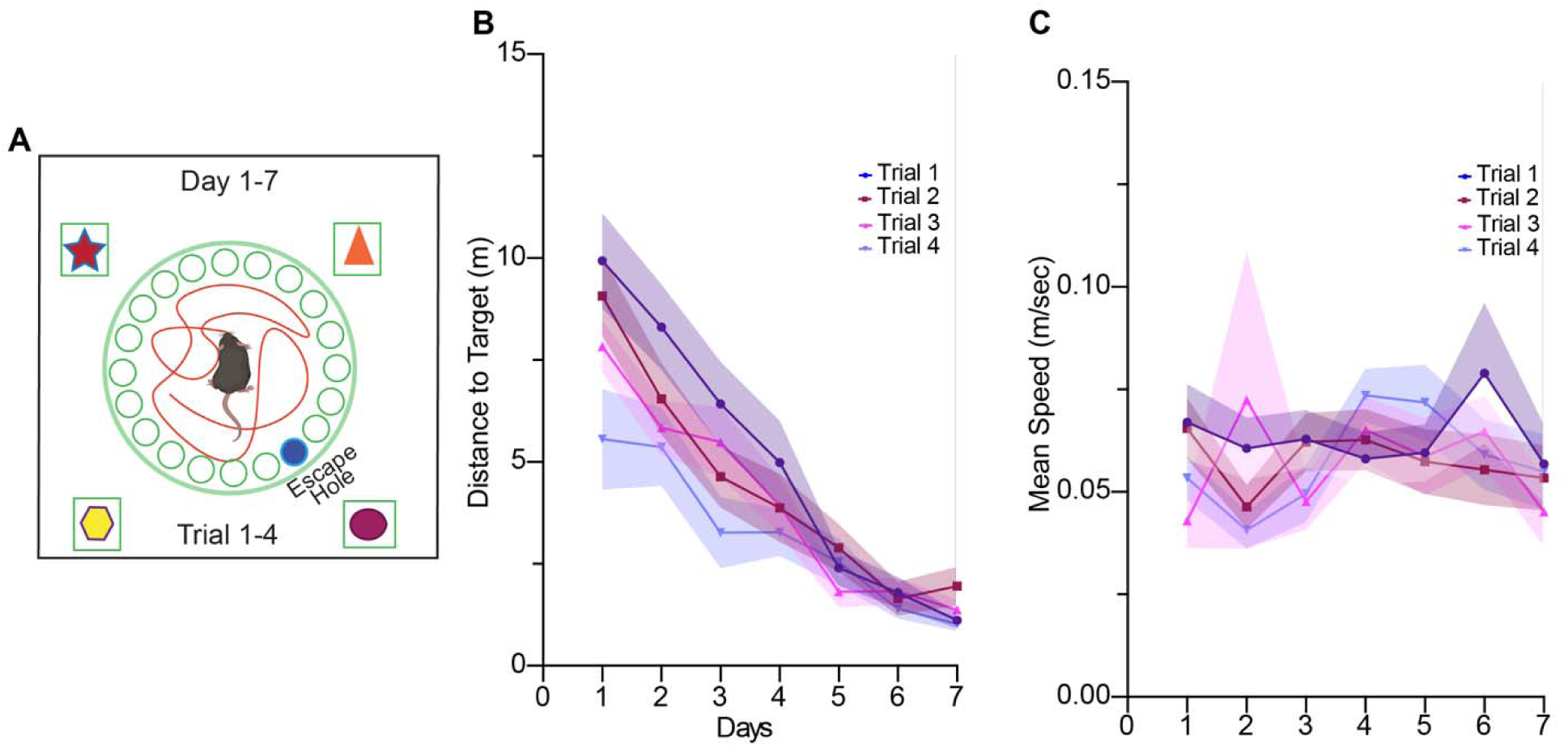
Distance and Mean speed to target hole. **(A)** Schematic diagram of the training for barnes maze with 4 trial per day. **(B)** Distance to target during learning sessions for 7 days in trial 1 (blue line), trial 2(dark red), trial 3(pink), trial 4(purple). For analyzing the differences between trials, statistical significance is fitting a full model two-way ANOVA, and no differences were observed among trials in each day. For analysis of differences across learning sessions days, statistical significance is an ordinary two-way ANOVA, confirming a significant effect between day 1 and day 2,3,4,5,6,7 (day 1 vs day 2, P= 0.0162, day 1 vs day 2,day 3,day 4,day 5,day 6,day 7, P<0.0001), day 2 and day 3,4,5,6,7 (day 2 vs day 3, P= 0.0069, day 2 vs day 4,5,6,7, P< 0.001), day 3 and day 5, 6, 7 (P<0.001) but not day 4, day 4 and day 5, 6, 7 (day 4 vs day 5, P= 0.0058, day 4 vs day 6,7, P<0.001) but not day 5 and day 6, 7. **(C)** Mean speed during learning sessions for 7 days in trial 1 (blue line), trial 2(dark red), trial 3(pink), trial 4(purple). For analyzing the differences between trials, statistical significance is fitting afull model two-way ANOVA, and no differences were observed among trials in each day. For analysis of differences across learning session days, statistical significance is fitting an ordinary two-way ANOVA, no differences were observed among learning days. Data are present as mean ± SEM.

## Acknowledgments

**To J.B.:** National Institutes of Health (NIH) National Institute of Neurological Disorders and Stroke (NINDS) BRAIN Initiative R01NS109994, NIH NINDS R01NS109362, NIH NINDS RM1NS132981, NIH National Institute of Mental Health (NIMH) R01MH122391, NIH National Institute on Aging (NIA) R01AG094086; Alzheimer’s Association AARGD-NTF-23-1151101, McKnight Scholar Award in Neuroscience, Klingenstein-Simons Fellowship Award in Neuroscience, Alfred P. Sloan Research Fellowship, Mathers Charitable Foundation Award, Whitehall Research Grant, a Blas Frangione Young Investigator Research Grant, New York University Whitehead Fellowship for Junior Faculty in Biomedical and Biological Sciences, and Leon Levy Foundation Award. **To M.H.-F.:** REC Scholar Program of NYU Alzheimer’s Disease Center (ADRC) P30 AG066512 and the Alzheimer’s Association grant 23AARFD-1026841. **To T.Z.:** NYU Dean’s Undergraduate Research Fund (DURF). **To C.D.J.:** NIH NIMH Diversity Supplement training award 3R01MH122391-04S1, parent award R01MH122391PIs Buzsáki/Basu. **To O.M.B.:** NIH 5T32NS86750 and REC Scholar Program of NYU Alzheimer’s Disease Center (ADRC) P30 AG066512.**To J.T.:** 2025 IES Brain Research Foundation Monique C. & George L. Braude Summer Fellowship in Neuroscience and the NYU College of Arts and Sciences Dean’s Undergraduate Research Fund (DURF).

We thank Drs. D. Lin, A. V. Masurkar. Hopkins and T. Butola for helpful discussion and inputs at various stages of the project and comments on the manuscript. We are grateful to Drs. V. Robert and J. Moore for input on the manuscript and support on the fiber photometry data analysis.

## Contributions

M. H.-F. and J.B. conceptualized and designed the experiments. J.B. secured primary funding, supervised, coordinated and managed the project. M. H.-F. performed all the experiments, analysis and prepared all the figures with input from J.B. and T.Z. T.Z. performed the scripts for the alignment of fiber photometry recordings with behavior and assisted with the analysis under supervision of M. H.-F. C.D.J. assisted with the chemogenetic experiments with training from M. H.-F. O. M. B. performed all the ex-vivo electrophysiological recordings and analysis. J.T. with supervision of M. H.-F assisted with immunohistochemistry and the anatomical analysis. M. H.-F. and J.B. wrote the paper with input from the authors.

